# Structure-based Discovery of Conformationally Selective Inhibitors of the Serotonin Transporter

**DOI:** 10.1101/2022.06.13.495991

**Authors:** Isha Singh, Anubha Seth, Christian B. Billesbølle, Joao Braz, Ramona M. Rodriguiz, Kasturi Roy, Bethlehem Bekele, Veronica Craik, Xi-Ping Huang, Danila Boytsov, Parnian Lak, Henry O’Donnell, Walter Sandtner, Bryan L. Roth, Allan I. Basbaum, William C. Wetsel, Aashish Manglik, Brian K. Shoichet, Gary Rudnick

**Author notes:** Co-corresponding give. Contributed equally (Isha Singh, Anubha Seth, Christian B. Billesbølle, Joao Braz).

## Abstract

The serotonin transporter (SERT) removes synaptic serotonin and is the target of anti-depressant drugs. SERT adopts three conformations: outward-open, occluded, and inward-open. All known inhibitors target the outward-open state except ibogaine, which has an unusual anti-depressant profile and stabilizes the inward-open conformation. Unfortunately, ibogaine is promiscuous and cardiotoxic, limiting understanding of inward-open state ligands. We computationally docked over 200 million small molecules against the ibogaine stabilized inward-open state of SERT. Thirty-six top-ranking compounds were synthesized and thirteen inhibited with potencies ranging from 29 to 5000 nM. Structure-based optimization led to two novel inhibitors with Ki values down to 3 nM. The new molecules stabilized an outward-closed state of the transporter and had little activity against off-targets. A cryo-EM structure of one of these bound to SERT confirmed the predicted geometry. In mouse behavioral assays, both had anxiolytic and anti-depressant activity, with potencies up to 200 better than fluoxetine.

## INTRODUCTION

The mammalian serotonin transporter (SERT) terminates the action of the neurotransmitter serotonin (5-hydroxytryptamine, 5-HT) released by serotonergic neurons. SERT catalyzes the transport of extracellular 5-HT back into the cell that released it, decreasing the 5-HT concentration in the synapse and recycling the neurotransmitter into the neuronal cytoplasm. There, it can be sequestered in synaptic vesicles for subsequent release (Rudnick and Clark, 1993). SERT catalyzed 5-HT transport requires Na^+^ and Cl^-^, as do transporters for other neurotransmitters such as norepinephrine (NE), dopamine (DA), γ-aminobutyric acid (GABA) and glycine (Gu et al., 1994; Guastella et al., 1990; Lingjaerde, 1969; Liu et al., 1992; Pacholczyk et al., 1991; Rudnick, 1977). All these neurotransmitter reuptake proteins belong to the NSS (neurotransmitter: sodium symporter), or *SLC6* (solute carrier 6) gene family, which also includes many animal and prokaryotic amino acid transporters that function outside the nervous system (Kristensen et al., 2011).

SERT (SLC6A4) is the target of many drugs, including antidepressants and psychostimulants. Competitive inhibitors selective for SERT are widely used therapeutically to treat clinical depression. Examples include fluoxetine, citalopram and paroxetine (Prozac, Celexa and Paxil) (Murphy et al., 2008). Cocaine, which also blocks the closely related transporters for NE and DA (NET and DAT), is a less selective competitive inhibitor for SERT (Gu *et al*., 1994). In addition, ibogaine, and its metabolite noribogaine, are alkaloids that non-competitively inhibit SERT and DAT (Bulling et al., 2012; Jacobs et al., 2007a) but are much less selective, with affinity for many receptors and channels (Glick et al., 2001). Distinct from these inhibitors, amphetamine derivatives, such as 3,4-methylenedioxymethamphetamine (MDMA, ecstasy) are substrates for SERT, NET and DAT that act, in part, by exchanging across the membrane with cytoplasmic neurotransmitters, thereby releasing transmitters into the synapse (Gu *et al*., 1994; Rudnick and Wall, 1992).

In the process of transporting 5-HT, SERT cycles through at least three distinct conformations. In the presence of extracellular Na^+^, SERT is predominantly in an outward-open conformation, in which an aqueous pathway connects the unoccupied orthosteric substrate site with the extracellular medium. Binding of extracellular 5-HT allows reversible conversion to an occluded state in which the substrate binding site is sealed off from the extracellular permeation pathway. In the presence of extracellular Na^+^, 5-HT and Cl^-^, the transporter reorients to an inward-open conformation in which a pathway opens between the substrate site and the cytoplasm releasing bound Na^+^ and 5-HT. Structures of SERT in these conformations (obtained by X-ray crystallography and by cryogenic electron microscopy (cryo-EM) (Coleman et al., 2016; Coleman et al., 2019; Yang and Gouaux, 2021)) align well to structures of other NSS transporters from both prokaryotes and animals (Krishnamurthy and Gouaux, 2012; Penmatsa et al., 2013; Shahsavar et al., 2021; Yamashita et al., 2005) indicating a common conformational mechanism within the family. In addition to these three conformations, an “inward occluded” conformation, observed in X-ray structures of the bacterial NSS transporters LeuT and MhsT, revealed a fourth state in which the extracellular pathway is closed as in the inward-open state, but the cytoplasmic pathway is also closed (Gotfryd et al., 2020; Malinauskaite et al., 2014).

In SERT and other NSS transporters, conformational changes open and close cytoplasmic and extracellular pathways to the central substrate site. The extracellular pathway opens and closes with the movement of a 4-helix bundle consisting of the extracellular halves of TMs 1, 2, 6 and 7 (Forrest et al., 2008) while the cytoplasmic pathway opens as the cytoplasmic half of TM1 splays out from its contacts with TM8 (Krishnamurthy and Gouaux, 2012). Separating the two processes allows transport to occur without substrate and ion leakage that might accompany the opening of one pathway simultaneously with closing the other.

The conformations revealed by X-ray and cryo-EM studies correlate with functional studies of NSS transporters using single molecule FRET, electron spin resonance and cysteine accessibility that demonstrate the extracellular pathway becoming more accessible in the presence of Na^+^ as the cytoplasmic pathway closes (Claxton et al., 2010; Fenollar-Ferrer et al., 2014; Forrest *et al*., 2008; Quick et al., 2006; Tavoulari et al., 2016; Zhang et al., 2021; Zhao et al., 2010). Addition of substrate, and of Cl^-^ in the case of mammalian neurotransmitter transporters, closes the extracellular pathway and opens the cytoplasmic pathway (Kazmier et al., 2014; Quick *et al*., 2006; Zhang *et al*., 2021; Zhang et al., 2018; Zhang et al., 2016; Zhao et al., 2011). SERT’s conformation also responds to the binding of antidepressants and of other drugs, with antidepressants and cocaine stabilizing the outward-open conformation while ibogaine and noribogaine stabilize the inward-open conformation (Coleman *et al*., 2019; Forrest *et al*., 2008; Jacobs *et al*., 2007a; Tavoulari et al., 2009).

The unique conformational properties of ibogaine and noribogaine lead to increased phosphorylation of SERT (Zhang *et al*., 2016) and raise the possibility that stabilizing the inward-open conformation could have unique pharmacological, and potentially therapeutic effects. For example, it has been reported that ibogaine ameliorates symptoms associated with opiate withdrawal (Alper et al., 1999; Brown, 2013; Schenberg et al., 2014; Wasko et al., 2018) and with depression (Glick *et al*., 2001; Rodriguez et al., 2020), although it is unknown whether these effects are SERT-mediated. Moreover, clinically relevant antidepressants stabilize SERT in outward-open conformations (Tavoulari *et al*., 2009), raising the question of how important is this conformational effect, as opposed to transport inhibition, in their antidepressant action. Unfortunately, ibogaine’s promiscuity toward other signaling receptors (Glick *et al*., 2001) has hindered studies that could more directly tie its conformational effect on SERT with any physiological outcome.

To investigate these questions, we sought compounds with effects on SERT akin to ibogaine but acting with greater selectivity. Structure-based docking of large chemical libraries has driven discovery of novel compounds that have high affinity and selectivity for several families of biological targets (Alon et al., 2021; Ballante et al., 2021; Bender et al., 2021; Gabrielsen et al., 2012; Gabrielsen et al., 2013; Gunera et al., 2020; Kampen et al., 2021; Katritch et al., 2012; Kolb et al., 2009a; Kolb and Irwin, 2009; Kufareva et al., 2012; Kufareva et al., 2017; Kufareva et al., 2014; Manglik et al., 2016; Ngo et al., 2016; Orry et al., 2006; Ortiz Zacarías et al., 2021; Patel et al., 2020; Rognan, 2012; 2017; Roth et al., 2017; Sadybekov et al., 2020; Scharf et al., 2019; Stauch et al., 2019; Stein et al., 2020; Uprety et al., 2021; Weiss et al., 2013; Wisler et al., 2018), though rarely transporters. We used the cryo-EM structure of SERT complexed with ibogaine in an inward-open conformation (Coleman *et al*., 2019) (PDB ID: 6DZZ) to computationally dock a library of 200 million make-on-demand molecules (Alon *et al*., 2021; Lyu et al., 2019; Sadybekov et al., 2022; Stein *et al*., 2020). A diverse set of high-ranking docked compounds, physically complementing the inward-open state of SERT and topologically unrelated to previously known inhibitors, were prioritized for synthesis and biochemical testing. From an original set of actives, mostly relatively potent, a cycle of structure-based design and testing led to two selective and potent compounds with anti-depressant and anxiolytic properties that act to close the extracellular pathway of SERT. A cryo-EM structure of one of these in complex with the transporter largely supports the docking-predicted geometry.

## RESULTS

### Retrospective control calculations

The recent determination of the cryo-EM structure of a SERT-ibogaine complex in an inward-open state (Coleman *et al*., 2019) afforded an opportunity to seek conformationally selective inhibitors. We targeted the extracellular-closed, inward-open state of the orthosteric site defined by residues such as Tyr95, Phe335 and Phe341. We further modeled two Na^+^ ions and one Cl^-^ ion that contribute to transport (Coleman *et al*., 2016; Rudnick, 1977) and for which sites are precedented in other transporter structures (Kantcheva et al., 2013; Penmatsa *et al*., 2013; Yamashita *et al*., 2005), though not explicitly seen in the ibogaine-SERT complex (Methods). We undertook retrospective control calculations against this site, seeking to confirm that we could preferentially dock ibogaine, noribogaine, 5-hyroxytryptamine (5-HT), cocaine, methylenedioxymethamphetamine (MDMA) and known selective serotonin reuptake inhibitors (SSRIs) (Tatsumi et al., 1997) in favorable geometries with high complementarity versus a set of property matched decoys (Mysinger et al., 2012); we find such control calculations to be helpful to optimize sampling and scoring parameters before undertaking a prospective docking campaign (Bender *et al*., 2021). We further investigated whether ibogaine and noribogaine would dock preferentially to the inward-open versus the other SERT conformations. Results of these retrospective calculations supported an ability to capture known conformationally-selective compounds in sensible geometries relative to property-matched decoys and to other states of the transporter (Figure S1).

### Ultra-large library docking versus the inward-open conformation of SERT

With the retrospective results in hand, we turned to docking a library of >200 million diverse, make-on demand molecules from the lead-like (Oprea, 2002) subset of ZINC (Irwin et al., 2020). These molecules, with molecular weights <350 amu, cLogP values < 3.5, among other restrictions, have favorable physical properties that allow for further optimization. Each library molecule was sampled for physical complementarity to the inward-open state of SERT by DOCK3.7 (Coleman *et al*., 2019). An average of 4358 orientations was sampled, and for each orientation about 187 conformations—over 1.57 x 10^11^ ligand configurations in total were sampled in 121,018 core hours (or 5 days over 1000 cores). High ranking molecules were filtered for interactions with Tyr95, Asp98, Tyr176, Ile172, Asn177, Phe335 and Phe341, those adopting strained conformations (Gu et al., 2021) were deprioritized, as were molecules that topologically resembled ∼28,000 annotated aminergic ligands acting at serotonin, dopamine and adrenergic receptors as well as known inhibitors of SERT, DAT or NET with ECFP4-based Tanimoto coefficients (Tcs) ≤0.35, based on molecules annotated in ChEMBL20 (Gaulton et al., 2017). Of the remaining molecules, the top ranking 300,000 were clustered for similarity to one-another. The best scoring members of 5000 of the resulting clusters (top 0.002% of the docked library) were visually inspected for engagement with critical residues in the orthosteric pocket of inward-open state of SERT, and for any new interactions in the binding pocket, using Chimera (Pettersen et al., 2004). These included salt bridge formation with Asp98, stacking with Phe335 and polar interactions with Asn177. Molecules with unsatisfied hydrogen bond donors and strained molecules were deprioritized. Ultimately, 49 molecules were selected for *de novo* synthesis and testing, out of which 36 were successfully synthesized (a 73.4% fulfillment rate). These 36 molecules are topologically both dissimilar to one another and dissimilar to known SERT inhibitors and complement the relatively unexplored inward-open state preferentially to the other two states seen to be adopted by SERT.

We first tested these 36 new compounds for SERT inhibition, beginning with the compounds at 30 μM (Figure 1A). Those molecules that inhibited greater than 50% of [^3^H]5-HT transport by SERT were considered active. Of the 36 tested, 13 molecules were active (≥ 50% inhibition, dashed line), a hit rate of 36% (hit rate = number-inhibited/number-physically-tested) (Figure 1A, Table S1). This relatively high hit rate is noteworthy as this docking campaign is among the first of which we are aware against a transporter. This supports the idea that structure-based docking is well-suited to targets outside of those families most heavily targeted until now, such as GPCRs, nuclear hormone receptors, kinases and soluble enzymes (Brown, 2013; de Graaf et al., 2011; Katritch et al., 2010; Kolb et al., 2009b; Mysinger *et al*., 2012; Powers et al., 2002; Wang et al., 2017).

**Figure 1.**
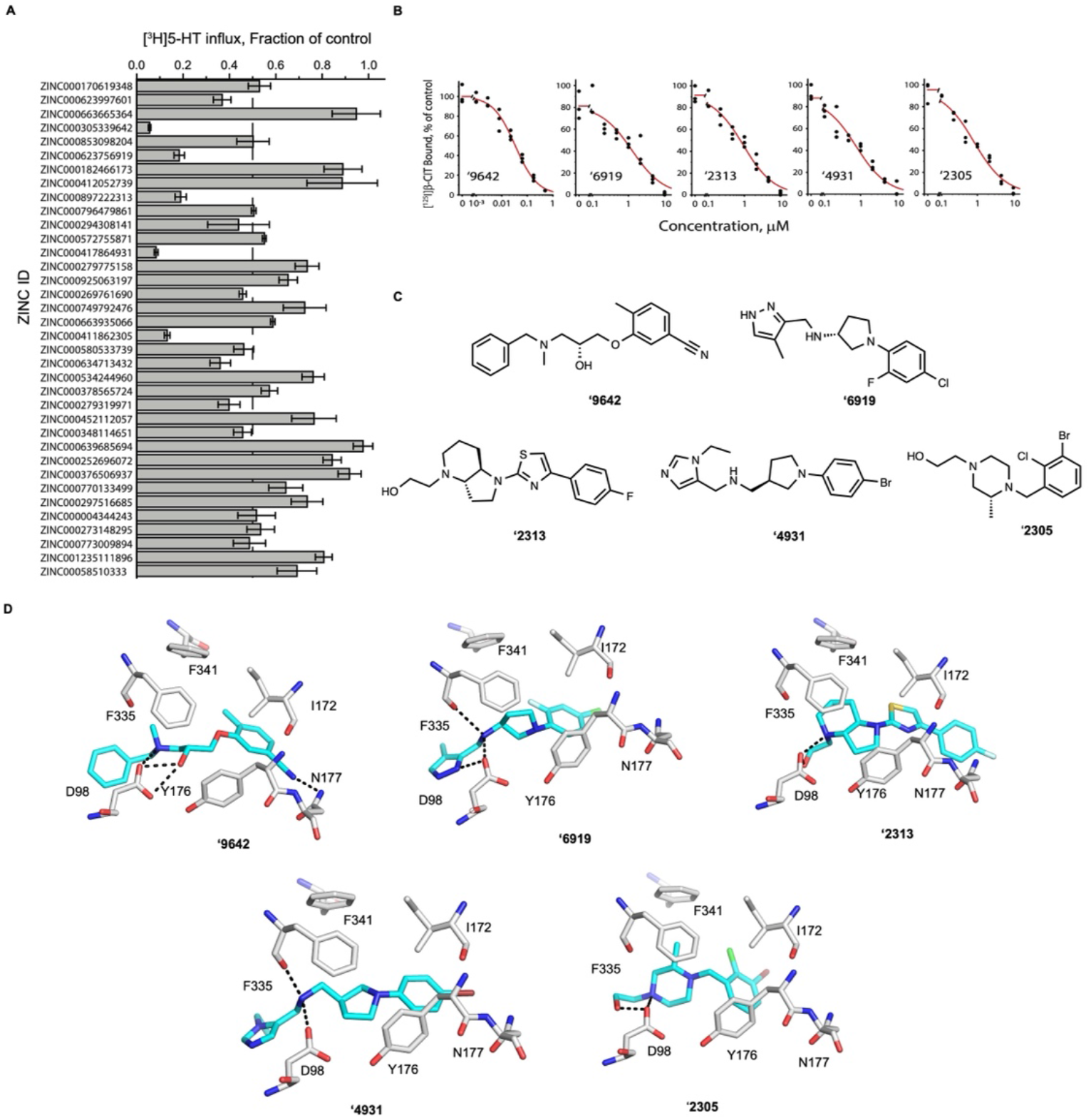
Novel inhibitors of serotonin transporter. (**A**) Inhibition of [^3^H]5-HT transport by molecules tested at 30 μM (mean ± SEM of three technical replicates). (**B**) Radio-ligand displacement curves of top five docking hits. (**C**) 2D Chemical structures of the top five docking hits; representing four different scaffolds. (D) Docked poses of top 5 hits including compounds ‘9642, ‘6919, ‘2313, ‘4931 and ‘2305. Protein ligand interactions are depicted as black dashed lines, ligands are represented as sticks with carbons in cyan and protein as sticks in with carbons in grey. Oxygens for both protein and ligand are in red, nitrogen in blue and sulfur in yellow.

To verify that the inhibition of transport represented interaction with SERT, we tested the five most potent hits (ZINC000305339642 or **‘9642**, ZINC000623756919 or **‘6919**, ZINC000897222313 or **‘2313**, ZINC000417864931 or **‘4931** and ZINC000411862305 or ‘**2305**) for their ability to displace the cocaine analog 2β-carbomethoxy-3β-(4-iodophenyl)tropane (β-CIT, RTI-55), a high-affinity SERT inhibitor. Figure 1B shows concentration-response curves confirming that these 5 docking hits were well-behaved inhibitors. These compounds inhibited [^125^I]β-CIT binding with affinities between 29 nM and 1.6 μM. The 2D chemical structures are represented in figure 1C. Docked poses of the five hits show unique interactions in SERT’s orthosteric site (Figure 1D).

### Structure guided optimization

We sought to improve the affinity of the five most active hits, representing 4 different novel chemotypes. Using the Smallworld (http://sw.docking.org) and Arthor (http://arthor.docking.org) search engines (NextMove Software, Cambridge UK) (Irwin *et al*., 2020), substructure and similarity searches were conducted among >20 billion make-on-demand Enamine REAL molecules, seeking analogs that well-complemented the SERT site. In a second approach, analogs that tested the particular modeled interactions, such as the salt bridge with Asp98, and the χ-χ stacking with Phe341 and Ph335, were bespoke synthesized; these were unavailable among the make-on-demand sets. Between 15 to 23 analogs were synthesized and tested for each scaffold (Table S2). Encouragingly, the overall affinities of the analogs increased by 2- to 700-fold over the parent molecules, and four of the five scaffolds saw an improvement in affinity (Figure S2). Most promising were analogs of ZINC000897222313 **(‘2313**), which itself had a Ki of 0.92 μM, with ZINC000006658090 (**‘8090**) and ZINC000443438219 (**‘8219,** (*R*)-enantiomer) having Ki values of 14 nM and 3 nM, respectively (Figure 2A). In the docked poses of the parent lead and its optimized analogs **‘2313**, ‘**8090** and ‘**8219**, modifications to the piperazine ring led to better stacking with Phe335 and Tyr176 (Figures 2B-D). Consistent with the specificity of these interactions, the (*S*)-enantiomer of ‘**8219**, ZINC000443438221 (**‘8221**), which differs only in the stereochemistry of a piperazine methyl, had a Ki of 170 nM, something reflected in the poorer docking pose adopted by **‘8221**, which does not make a favorable hydrogen bond to the recognition Asp98, owing to packing flaws introduced by the methyl stereochemistry (Figure S3).

**Figure 2.**
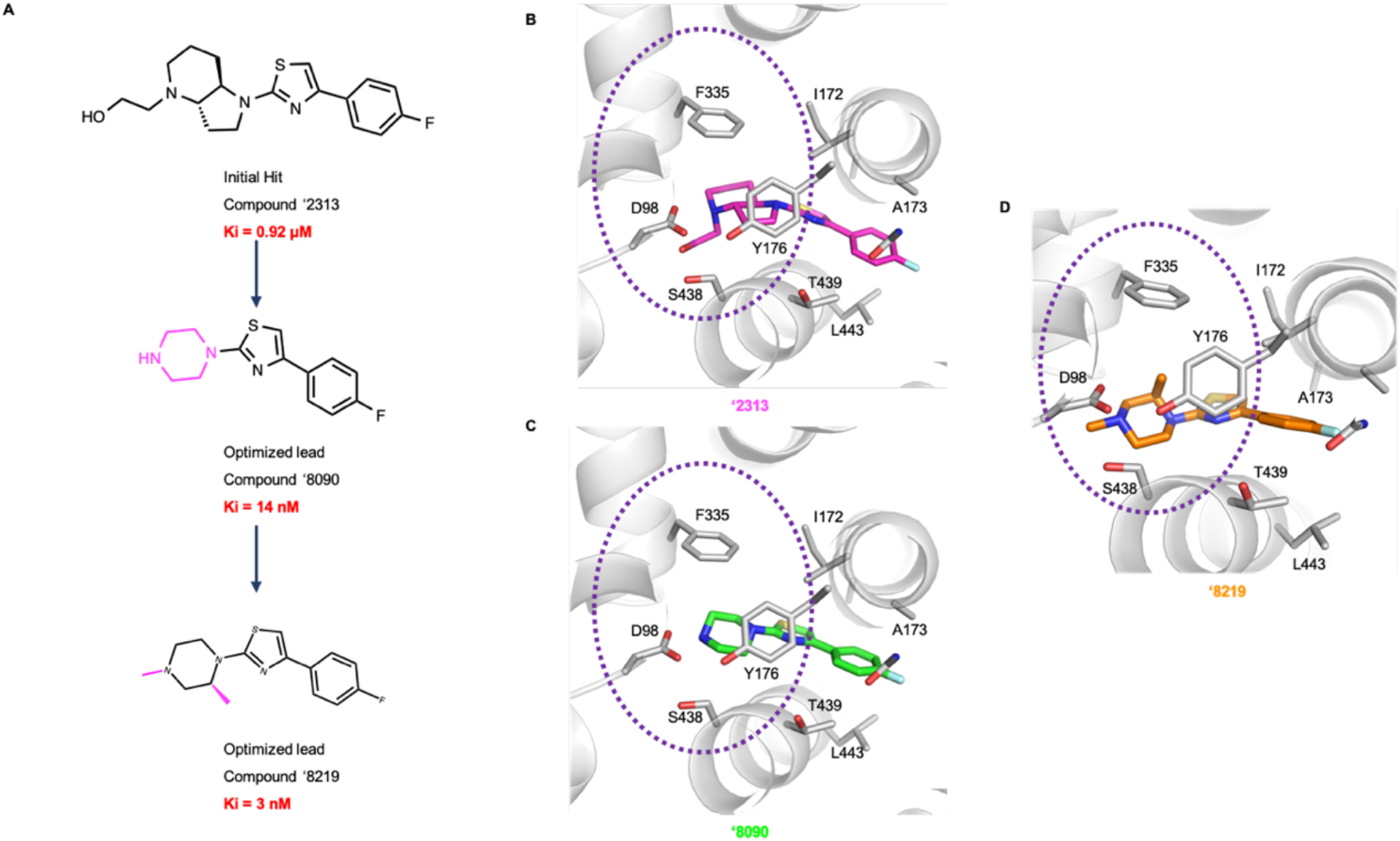
Hit to lead optimization of the compound ‘2313 leading to nM SERT inhibitors. (A) Chemical structures of the parent compound, **‘2313** and its corresponding analogs including compound **‘8090** (Ki = 14 nM) and compound **‘8219** (Ki = 3 nM). The variable group in the analogs with respect to the parent compound are colored in magenta. (B) Docked pose of the parent compound, **‘2313**, (C) the optimized lead **‘8090** (D) and the most potent lead **‘8219** are represented as cartoon. Dashed circles represent the improved stacking of F335 with ring substitutions in going from parent compound to the most potent lead.

### Molecular interactions of ‘8090 and ‘8219 with SERT

We wished to investigate the kinetic mechanism of inhibition of the new inhibitors, and the conformation of SERT to which they preferentially bound. In particular, we were interested to know whether the new ligands were competitive substrates or true inhibitors, and the conformation of the SERT to which they preferentially bound (recall that we were initially motivated by seeking ligands that captured SERT selectively in the rare inward-open state). To address these questions, we undertook both straightforward competition experiments, probed the dependence of inhibition on sodium and chloride ion concentrations, which are known to preferentially stabilize different transporter states, and explored the exposure of residues along the extra- and intra-cellular pathways in response to inhibitor binding, using cysteine modification assays.

#### **‘8090** and **‘8219** are not substrates

Binding and transport studies show that these compounds are not SERT substrates, but rather non-competitive inhibitors like ibogaine. Both **‘8090** and **‘8219** decreased the Vmax for 5-HT influx with minimal effect on K_M_, indicating non-competitive inhibition (Figure 3A), an observation suggesting that they do not interfere with 5-HT binding from the external medium and therefore are not substrates. To test this directly, we exploited an exchange assay to measure the ability of SERT to exchange accumulated [^3^H]5-HT with extracellular compounds. Extracellular unlabeled 5-HT, as a substrate, stimulated robust efflux of previously accumulated intracellular [^3^H]5-HT as previously described (Rudnick and Wall, 1992; Wall et al., 1995) (Figure 3B). However, neither **‘8090** nor **‘8219** significantly increased efflux when added at saturating concentrations, suggesting that they were not SERT substrates.

**Figure 3.**
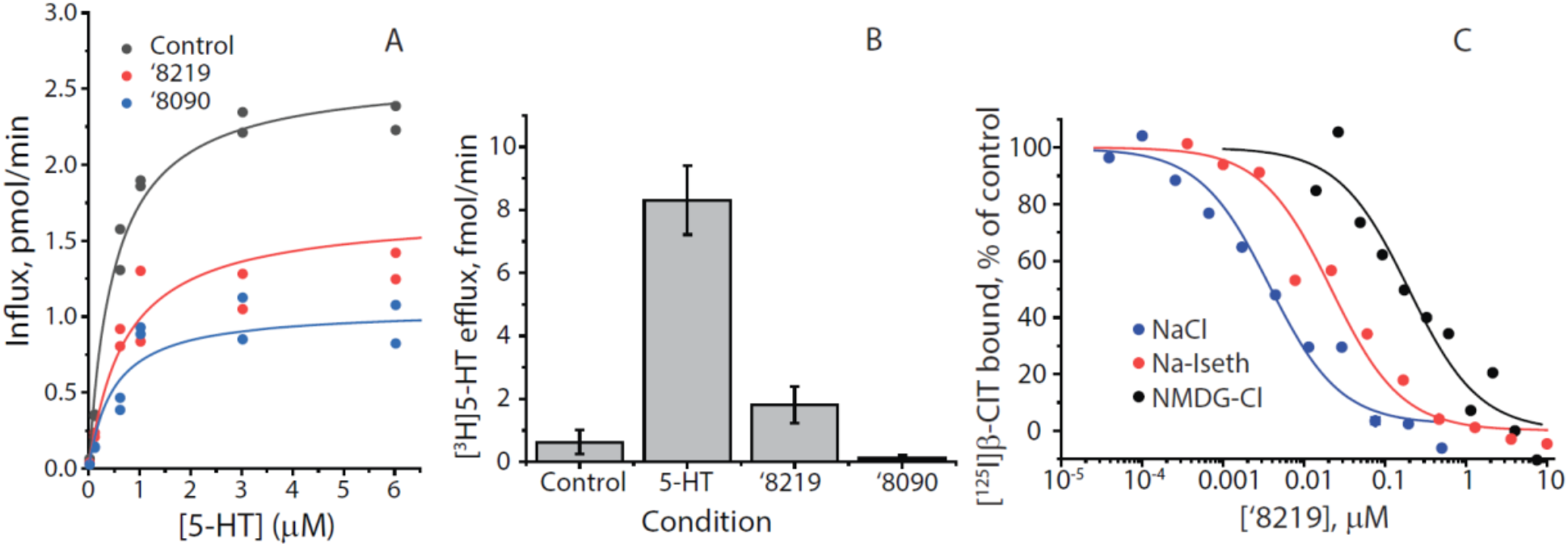
Molecular interactions of SERT with ‘8219 and ‘8090. **(A)** Kinetics of inhibition by 0.15 μM **‘8219** and 0.7 μM **‘8090** (representative experiment). 5-HT transport into HeLa cells transfected with hSERT was non-competitively inhibited by both compounds. There was a small but statistically insignificant increase in K_M_ with **‘8219** (0.69 ± 0.25 μM (SEM), n=6) and a decrease with **‘8090** (0.47 ± 0.05 μM, n=5) compared with uninhibited control (0.59 ± 0.18 μM, n=8). P values for paired t-tests were 0.99 and 0.18, respectively. Vmax was significantly decreased for both compounds from 1.7 ± 0.25 pmol/m/well for control (n=8) to 1.1 ± 0.2 pmol/m/well (n=6) for **‘8219** and 0.69 ± 0.1 pmol/m/well (n=5) for **‘8090**. P values for paired t-tests were 0.018 and 0.011, respectively. **(B)** Efflux of accumulated [^3^H]5-HT induced by extracellular unlabeled 20 μM 5-HT, 10 μM **‘8219** or 10 μM **‘8090**. 5-HT induced marked efflux of radiolabel, 8.3 ± 1.1 fmol/m (SEM), relative to control (no addition) 0.63 ± 0.39 fmol/m (P=7×10^-4^ in 2-sample t-test, n=6). **‘8219** slightly increased efflux but the increase was not significant (P=0.18) and **‘8219** did not increase efflux. **(C)** Na^+^ and Cl^-^ increased **‘8219** affinity in equilibrium displacement of [^125^I]β-CIT. Membranes from cells expressing SERT were incubated with 0.1 nM [^125^I]β-CIT and the indicated concentrations of **‘8219** in PBS (control), PBS in which Na^+^ was replaced with NMDG^+^ (black line and circles) or Cl^-^ was replaced with isethionate (red line and circles). The presence of Cl^-^ increased **‘8219** inhibitory potency 9-fold, from a K_I_ of 37 ± 11 nM to 4.0 ± 0.4 nM in these experiments (n-3). Na^+^ increased **‘8219** inhibitory potency 127-fold, from a K_I_ of 508 ± 56 nM to 4.0 ± 0.4 nM

#### Affinity depends on cellular context

Much higher concentrations of these compounds were required to inhibit 5-HT influx into intact cells versus inhibition of β-CIT binding to membrane preparations (Figure S4A). Although this difference could have resulted from slow binding kinetics (90 min equilibrium binding measurement vs. 10 min transport assay) preincubation with either compound for 30 min did not decrease the K_I_ for transport (Fig. S5A, Tables S4, S6). We also measured the dissociation rate of **‘8090** and **‘8219** by following the recovery of SERT-mediated 5-HT-dependent ionic currents. SERT expressing cells were saturated with either **‘8090** or **‘8219** and the recovery of a 5-HT current was measured during washout (Figure S4B). On-rates calculated from these dissociation rates and measured K_D_ values were fast enough that the compounds would completely equilibrate with SERT during the 30-minute pre-incubation (Table S5). The difference in potency was likely due to the cellular vs membrane context of SERT, possibly because the compounds access the binding site from the cytoplasmic side.

#### The effects of sodium and chloride ions on binding

The compounds differ from ibogaine in their unique ionic requirements and effects on conformation. The ion dependence of **‘8090** and **‘8219** binding, measured by displacement of [^125^I]β-CIT, differed dramatically from that of ibogaine. Ibogaine binding to SERT was *inhibited* by Na^+^ and independent of Cl^-^ (Tavoulari *et al*., 2009). In contrast, binding of **‘8219** was *enhanced* over 120-fold by Na^+^ over 9-fold by Cl^-^ (Figure 3C). For **‘8090**, these effects were smaller, with a 10-fold enhancement for Na^+^ and a statistically insignificant effect of Cl^-^ (Figure S5 and Table S6).

#### Effect on SERT conformation

Because Na^+^ and Cl^-^ influence the conformation of SERT and related transporters (Zhang *et al*., 2021; Zhang *et al*., 2016), we tested the effect of **‘8090** and **‘8219** on SERT conformation, seeking to understand if the new inhibitors bound to the inward-open conformation, as targeted, and akin to ibogaine, the outward open conformation, akin to cocaine and SSRIs, or an intermediate state. We used SERT mutants, depleted in reactive endogenous cysteine residues, and containing novel cysteine residues in either cytoplasmic or extracellular pathways, as previously used to determine the conformational effects of ibogaine on SERT (Jacobs et al., 2007b) (see Figure S6). Figure 4A shows how the reactivity of a cysteine replacing Ser277 in the cytoplasmic pathway is affected by the five top hits, **‘8090** and **‘8219**, and several reference compounds. As previously reported, cocaine and ibogaine have opposing effects on the accessibility of this residue (Jacobs *et al*., 2007b). Cocaine, by stabilizing SERT in an outward-open conformation, *decreased* Cys277 reactivity, relative to control (no addition, dashed line), because the cytoplasmic pathway is closed in this conformation (Forrest *et al*., 2008). In contrast, ibogaine, by stabilizing SERT in an inward-open conformation, dramatically *increased* Cys277 reactivity.

**Figure 4.**
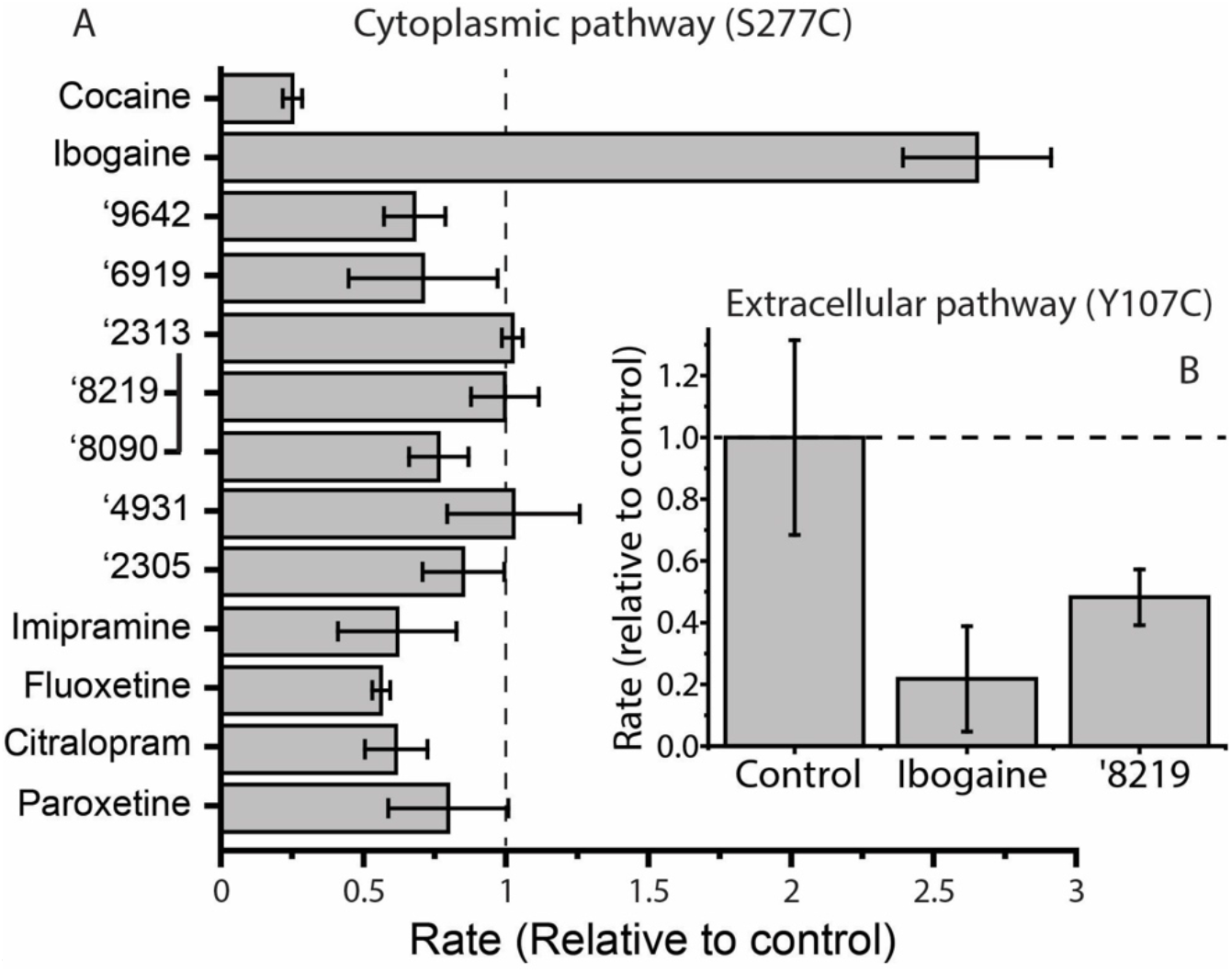
Influence of ‘8219 and ibogaine on SERT conformation. **(A)** Effects on Cys277 reactivity in the cytoplasmic pathway. Rates of Cys277 modification in membranes from HeLa cells expressing SERT-S277C by MTSEA was measured by the decrease in β-CIT binding activity after treatment with MTSEA in the presence of the indicated compounds, each present at a saturating concentration (10x K_D_). The reference compounds cocaine and ibogaine are shown at the top, and several clinically used antidepressants are shown at the bottom of the plot. Between them are the five top hits shown in Figure 1 and the two analogs of **‘2313** which were the highest affinity compounds found in this project, **‘8219** and **‘8090**. The control rate was 75 ± 15 s^-1^M^-1^. **(B)** Effects on Cys107 reactivity in the extracellular pathway. Rates of Cys107 modification in HeLa cells expressing SERT-Y107C was measured by the decrease in residual transport activity after treatment with MTSET in the presence of the indicated compounds, each present at a saturating concentration (10x K_D_). The control rate was 217 ± 68 s^-1^M^-1^.

To our surprise, none of the compounds that emerged from this screen mimicked ibogaine by increasing Cys277 reactivity relative to control. Several, including **‘8090**, decreased reactivity slightly, but not nearly as much as cocaine. In general, the effects of these compounds on the cytoplasmic pathway were similar to those of commonly used antidepressant compounds, and **‘8219** seemed not to affect cytoplasmic pathway at all.

To compliment these studies, we measured the effect of **‘8219** on the reactivity of a cysteine replacing Tyr107 in the *extracellular* pathway. Cys107 in this SERT mutant reacts in the opposite way to Cys277 in the presence of cocaine and ibogaine; cocaine renders it more accessible and ibogaine decreases its accessibility (Jacobs et al, 2007). Here, compound **‘8219** acted to decrease Cys107 reactivity as ibogaine does and unlike SSRIs, which open the extracellular pathway (Tavoulari *et al*., 2009). This even though it had little or no effect on the cytoplasmic pathway.

Returning to our motivating questions, we find that the new inhibitors, **‘8090** and **‘8219**, are non-competitive inhibitors and not substrates, akin to ibogaine. Unlike ibogaine, they do not preferentially stabilize an open cytoplasmic pathway but like ibogaine and *unlike* classic SSRIs they decrease the accessibility of the extracellular pathway. Taken together, these results suggest that the primary effect of these new SERT inhibitors is to close the extracellular pathway, with little or no effect on the cytoplasmic pathway.

### Cryo-EM structure of a SERT/’8090 co-complex

To test our docked model, we sought experimental structures of our optimized analogs in complex with SERT. Our attempts to obtain structures of SERT in the presence of K^+^, which stabilizes the inward-open state used in our docking campaigns, were unsuccessful. We were, however, able to determine a 3.0 Å structure of nanodisc-reconstituted (Nasr et al., 2017) SERT bound to Fab15B8 (Coleman *et al*., 2019) and compound ‘**8090** in the presence of Na^+^ (PDB: 7TXT, EMBD: EMD-26160, Figure 5A and S7). The resulting map yielded well-defined and contiguous TM densities that allowed for unambiguous modeling of SERT regions important for ligand binding (Figure S7). A non-proteinaceous density overlapping with the orthosteric site, and with a local resolution of ∼ 2.7 Å, allowed for confident modeling of **‘8090** (Figure 5A and Figure S7).

**Figure 5.**
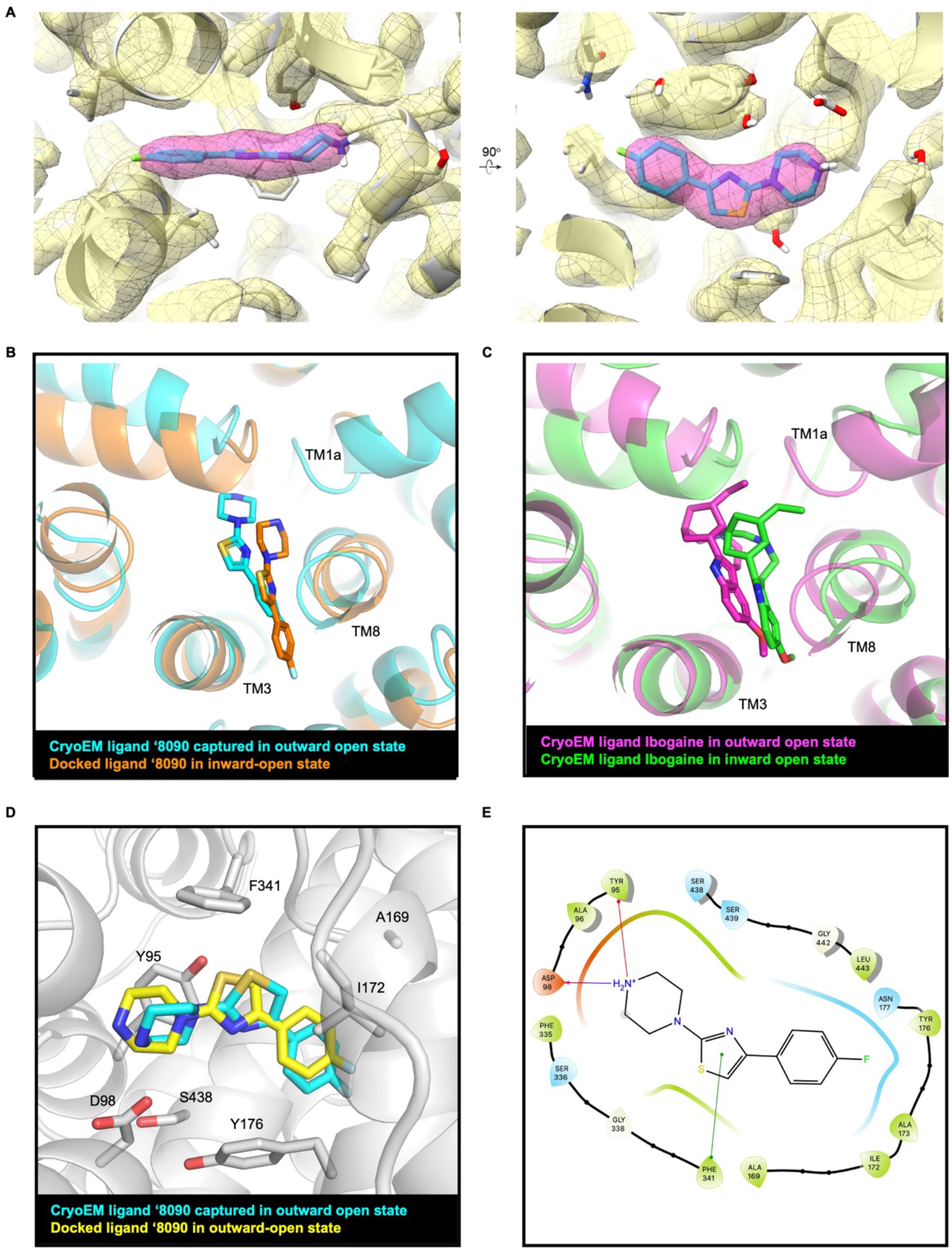
Structural fidelity between docked and cryo-EM determined pose of ‘8090. (**A**) Cryo-EM density maps of compound ‘**8090** bound to SERT in the outward-open state (PDB: 7TXT, EMBD: EMD-26160. (**B**) Comparison of **‘8090** binding pose in cryo-EM determined structure (cyan) and respective docked pose in the inward open state of SERT (orange). (**C**) Comparison of ibogaine binding pose in cryo-EM determined inward open state structure (green, PDB ID: 6DZZ) and outward open state of structure of SERT (magenta, PDB ID: 6DZY). (**D**) Ligand cryo-EM determined pose (cyan) overlaid with respective docked pose (yellow) in the outward open active site of SERT. (**E**) 2D outline of protein−ligand interactions for **‘8090** interacting with SERT.

While Na^+^ dramatically increased the affinity of **‘8219** and related compounds (Fig. 3D, S4), our *in vitro* accessibility assays indicated that **‘8219** favors SERT in an outward-closed state (Fig. 4B). Taken together, our in vitro data could reflect an equilibrium between open and closed states of the extracellular pathway induced by **‘8090**, similar to that reported for 5-HT bound SERT in Na^+^ (Yang and Gouaux, 2021). However, exhaustive processing converged on just a single structure of ‘**8090**-bound SERT in an outward-open conformation (1.43Å all-atom RMSD relative to paroxetine-bound SERT in an outward-open conformation, PDB ID:6VRH (Coleman *et al*., 2019)), with extracellular gating residues (Arg104 and Glu493; Tyr176 and Phe335) oriented to accommodate an extracellular solvent pathway reaching into the orthosteric site.

It should be noted that cryo-EM structures of ibogaine bound to detergent-solubilized SERT in the presence of Na^+^ (Coleman *et al*., 2019) show the transporter to be in an inward-closed conformation despite accessibility measurements of SERT in native membranes that show that ibogaine dramatically *opens* the cytoplasmic pathway and *closes* the extracellular pathway under the same conditions (Jacobs *et al*., 2007a) (Fig. 4). The reasons for this discrepancy are not clear, but it implies that the cryo-EM data may not capture the full dynamic range of SERT conformation under these conditions.

Because the architecture of the SERT orthosteric site is highly similar among conformational states of the catalytic transport cycle, we nevertheless reasoned that our structure might provide insight into the ‘**8090** binding pose in other conformations. On comparing the cryo-EM predicted binding pose of **‘8090** bound in the outward-open state of the transporter to the docked pose of **‘8090** in the inward-open state to which the ultra-large scale docking was carried out, they aligned in the same orientation with fluorine protruding into the cavity between TM3 and TM8 between the amino acid residues Asn177 and Ile172 (Figures 5B and 5D-E). Similarly, cryo-EM structures of ibogaine obtained in all three SERT conformations show a conserved pose of ibogaine with the methoxy group of ibogaine protruding into the cavity between TM3 and TM8 (Figure 5C). As the transporter transitions from outward-open state to inward-open state, the position of the ligand is adjusted, moving it in the direction of TM1a and TM8, depicted here in Figure 5B for **‘8090** and 5C for ibogaine.

The experimental maps for **‘8090** broadly supports its docked binding pose, which occupies the SERT orthosteric site with key interacting residues Asp98, Tyr95 and Phe341, akin to the docking prediction (Figure 5D-E). To compare like-to-like structures, **‘8090** was re-docked to the outward open conformation (PDB: 6DZY). In this transporter conformation, the docked and cryo EM determined ligand superposed even more closely with the EM map, with an RMSD value of 1.17 Å (Figure 5D, Supplementary Table S3 and Figure S7). In both the docked binding poses and cryo-EM fitted ligand, the compound ‘**8090** stacks with Phe341 and forms salt bridge with Asp98.

### Both ‘8090 and ‘8219 are selective versus NET, DAT, and ∼300 GPCR targets

To compare inhibitory activity at neurotransmitter transporters, we tested compounds ‘**8090** and ‘**8219** in cell-based functional assays. Against DAT and NET ‘**8090** had Ki values of 6.6 and >10,000 μM, respectively, while ‘**8219** had Ki values of 7 and 3.5 μM, respectively. (Figure S8). Both ‘**8090** and ‘**8219** were also tested for off-target agonist activity at > 300 human GPCRs in the Tango assay (Kroeze et al., 2015) at a concentration of 10 µM; little activity was seen against any target except for the 5-HT_1A_ receptor. In concentration-response Gi activity assays, ‘**8090** had weak Gi agonist activity at 5-HT_1A_, with about 6% activity of reference agonist 5-HT (Figure S8).

### The new SERT inhibitors have anti-anxiolytic and anti-depressant activities in mice

Dysregulation of SERT has been linked to psychiatric disorders such as major depressive disorder (MDD), and SERT is the therapeutic target of well-known anti-depressants such as fluoxetine, citalopram, and paroxetine. With their ability to close the extracellular pathway, which differs from the action of classic SSRIs, it seemed interesting to test compounds **‘8090** and **‘8219** for anti-anxiolytic and anti-depressant activity. The two molecules were first tested for pharmacokinetic exposure. On intraperitoneal (IP) dosing, **‘8219** and **‘8090** achieved brain C_max_ values of 3530 ng/g and 12000 ng/g with half-lives of 147 and 44.3 minutes, respectively (Figure S9). These relatively high exposure levels reflect the favorable physical properties for which these molecules were selected and optimized: both are cations with molecular weights < 300 amu with low topological polar surface area, well-suited for exposure in the CNS.

Bolstered by the pharmacokinetics, the two inhibitors were tested in two mouse behavioral assays: the elevated plus-maze (thought to map to anxiety), and the acute and 24 h tail suspension assay (thought to map to depression). In the test for anxiety-like behavior, both ‘**8219** and ‘**8090** were chronically injected in mice for 10 days at concentrations of 10 mg/kg. The SSRI paroxetine (10 mg/kg) was used as a positive control. Compared to vehicle, both of the new compounds increased the percentage of entries into and the time spent exploring the open arms (Figure 6A). Conversely, the time spent in the closed arms decreased for both compounds. Importantly, the number of total entries as well as the total distance travelled in the plus maze apparatus over the 5 min testing period were not significantly different between the groups (Figure 6B), indicating that the compounds did not alter the motor performance and/or exploratory behavior of the mice. As expected, mice receiving paroxetine also entered and spent more time exploring the open arms, compared to vehicle injected mice (Figure 6C,D). Thus, both compounds ‘**8219** and ‘**8090** produced an anti-anxiolytic-like effect in the plus-maze test.

**Figure 6.**
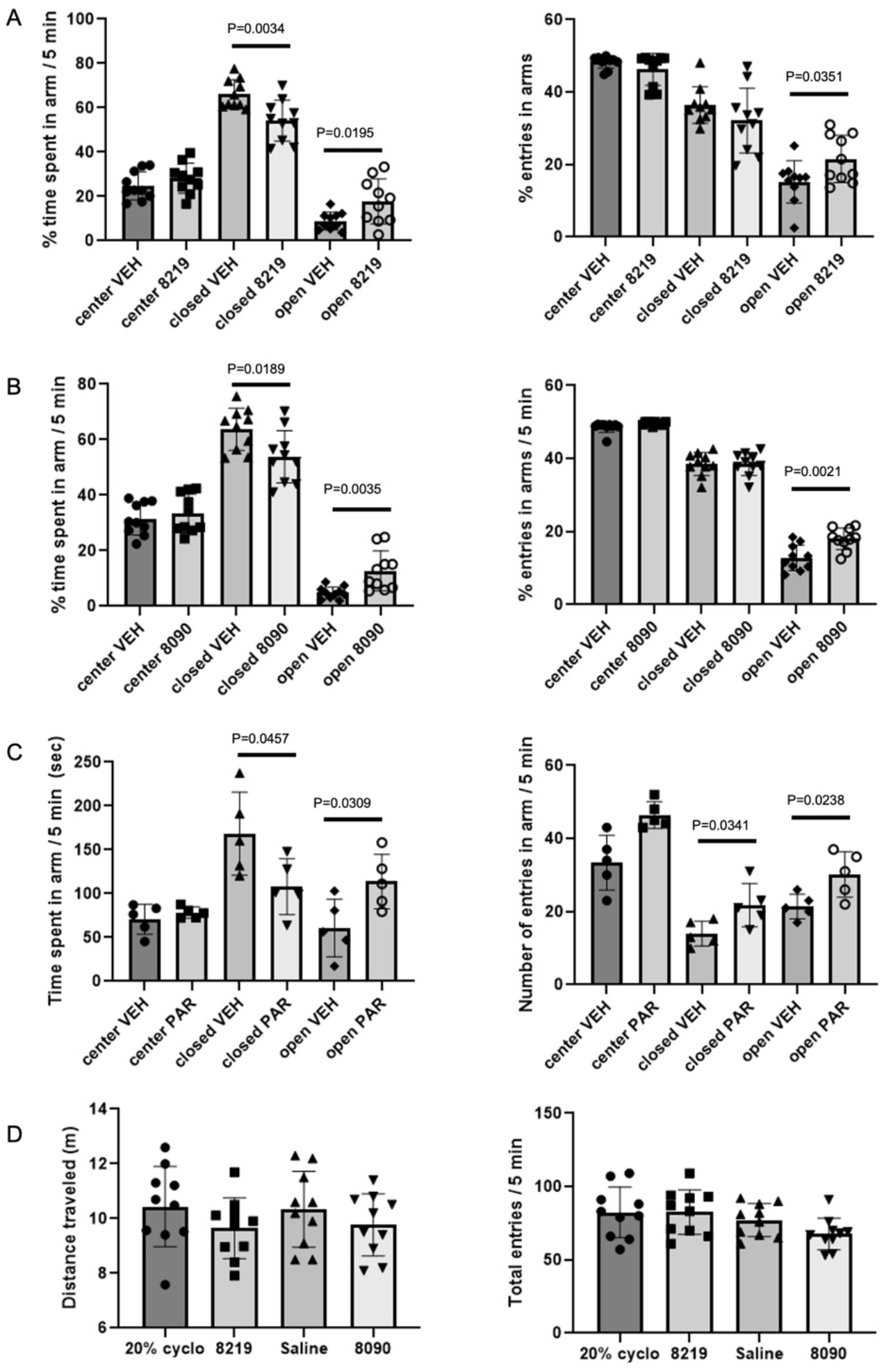
Anxiolytic effects of compounds 8219 and 8090 in the Elevated plus maze assay. **(A-B)** Effect of compounds **8219** (A; 10 mg/kg for 10 days; N=10) and **8090** (B; 10 mg/kg for 10 days; N=10) on the time spent (left) and the number of entries (right) into each arm (center, closed and open) of the elevated plus maze. **(C)** Effect of a single administration of paroxetine (10 mg/kg; N=5) on the time spent (left) and the number of entries (right) into each arm (center, closed and open) of the elevated plus maze. **(D)** Effect of compounds, **8219** (N=10) and **8090** (N=10) on the traveled distance (left) and the total number of entries (right) in the elevated plus maze. Data are presented as mean ± SEM. Significance levels were determined using an unpaired Student’s t-test and comparing the effect of compounds **8219** and **8090** to their own vehicle (20% cyclodextrin and saline respectively).

Both inhibitors were then tested in a tail suspension assay for anti-depressant-like activity 30 min and 24 h after injection. For this experiment, VMAT2 mice were used. Parenthetically, chronic inhibition of the VMAT with reserpine in human hypertensive patients produced depression without anxiety (Freis, 1954) and this effect provided a basis for the monoamine hypothesis of depression (Maes et al., 1994; Schildkraut, 1965). Homozygous deletion of *Vmat2* is lethal in mice, while VMAT2 heterozygous (HET) mice survive and are healthy (Wang et al., 1997). The adult male and female VMAT2 HET mice are hypoactive in the open field, display anhedonia-like behavior with sucrose solutions, and show increased immobility in the forced swim and tail suspension tests which were alleviated with tricyclic anti-depressants, SSRIs, selective norepinephrine transporter inhibitors, and the atypical anti-depressant bupropion (Fukui et al., 2007). These mutants do not present with anxiety-like responses, but display enhanced learned helplessness and increased responses to stress compared to WT controls. The well-known SSRI fluoxetine (Prozac) was used as a positive control in the present experiment.

As expected, immobility in the tail suspension test was high both acutely (30 min after injection), as well as 24 h later in the vehicle-treated VMAT2 HETs versus their wild-type (WT) controls (p-values<0.001) (Figure 7A-D). After acute dosing, all doses of ‘**8090** and ‘**8219** reduced immobility versus vehicle, with the exception of 0.5 mg/kg ‘**8090** (Figure 7A). While 20 mg/kg fluoxetine also substantially reduced immobility, it did so at at concentrations 40-fold to 200-fold higher than equi-efficacious doses of **‘8090** and ‘**8219**, respectively. Even at 24 h post-dose, anti-depressant-like activities were maintained with 0.5 and 2 mg/kg ‘**8090** and with all doses of ‘**8219** versus the vehicle control for the VMAT2 HET mice (Figure 7B), while 2 mg/kg ‘**8090**, and 0.5 and 2 mg/kg ‘**8219** were more efficacious than fluoxetine. In comparisons of responses in VMAT2 HETs across time, antidepressant activities were maintained across 24 h with fluoxetine and both compounds at all doses, except for 1 mg/kg ‘**8090** (Figure 7A-B). In summary, vehicle-treated VMAT2 HET mice displayed depressive-like behaviors relative to WT controls at 30 min and 24 h after injection. Both ‘**8090** and ‘**8219** had potent anti-depressive-like actions, up to 200-fold more potent than fluoxetine, and these effects were maintained over 24 h. These results are intriguing considering the relatively equal potency of the new compounds with fluoxetine in inhibiting SERT function in vitro.

**Figure 7:**
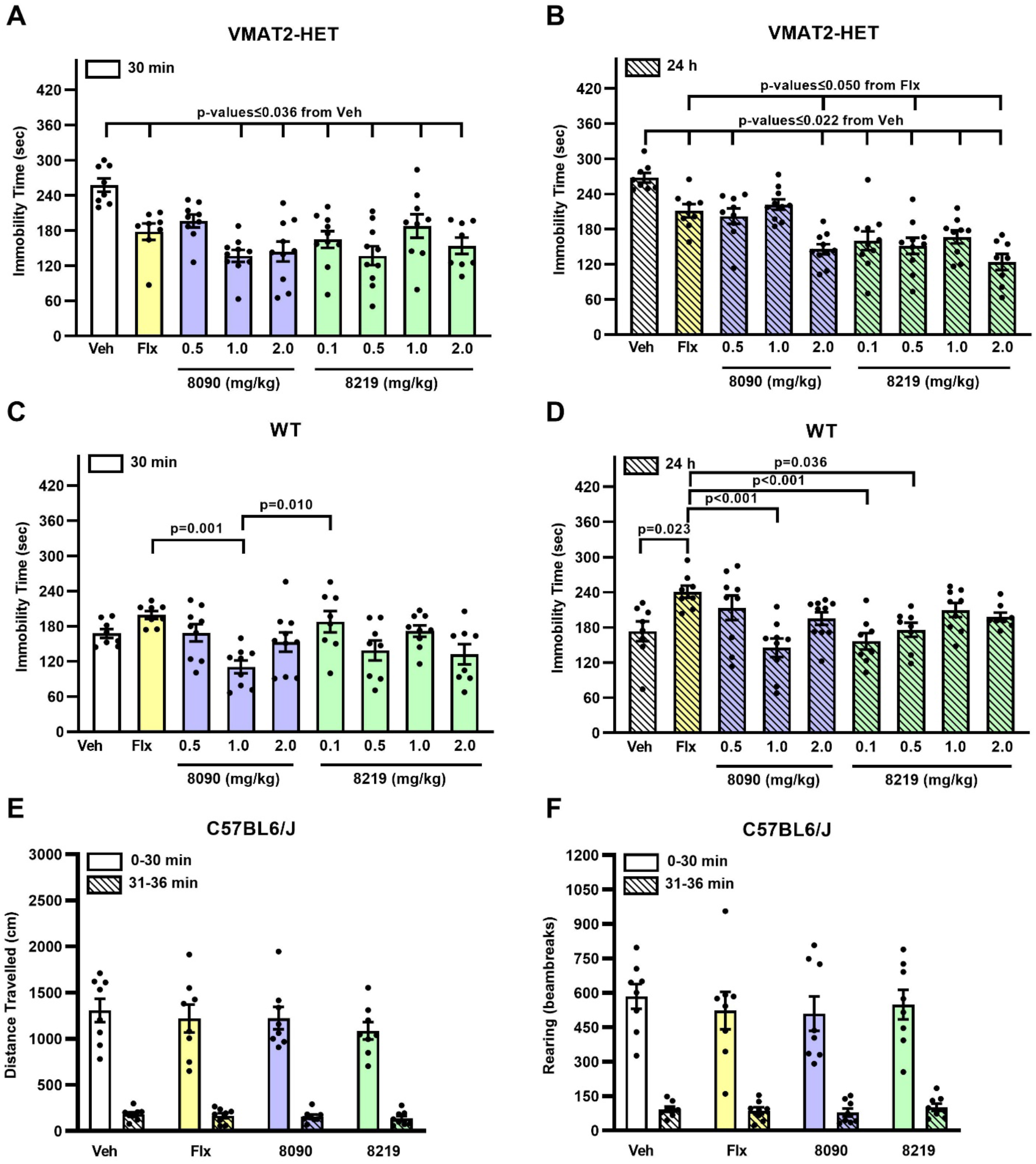
Anti-depressive-like effects of compounds 8090 and 8219 in the 2-day tail suspension test with *Vmat2* mice. **(A-B)** Effects of compounds ‘**8090** (0.5, 1, and 2 mg/kg, i.p.) and ‘**8219** (0.1, 0.5, 1, and 2 mg/kg, i.p.) at 30 min (acute) 24 h after injection in VMAT2 heterozygous (HET) mice. Controls were administered (i.p.) the vehicle (Veh) or 20 mg/kg fluoxetine (Flx). **(C-D)** Effects of compounds ‘**8090** (0.5, 1, and 2 mg/kg, i.p.) and ‘**8219** (0.1, 0.5, 1, and 2 mg/kg, i.p.) at 30 min and 24 hr after injection in wild-type (WT) mice. RMANOVA within subjects tests for time [F(1,141)=20.754, p<0.001], time x genotype [F(1,141)=5.032, p=0.026], time x treatment [F(8,141)=2.557, p=0.012], time x genotype x treatment [F(8,141)=2.677, p=0.009], and the between subjects tests for treatment [F(8,141)=12.831, p<0.001] and genotype x treatment [F(8,141)=8.557, p<0.001]. Bonferroni corrected pairwise comparisons, p<0.05. N=8-10 mice/genotype/treatment/time. **(E-F)** Effects of compounds ‘**8090** (0.5, 1, and 2 mg/kg, i.p.) and ‘**8219** (0.1, 0.5, 1, and 2 mg/kg, i.p.) at 0-30 min and at 31-36 min (corresponding to time of testing tail suspension) after injection in C57BL/6J mice on motor activities in the open field. Controls were administered (i.p.) the vehicle (Veh) or 20 mg/kg fluoxetine (Flx). RMANOVA identified no significant effects. N=8 mice/treatment.

Since drugs/compounds that increase motor activities can confound the results from the tail suspension test (Steru et al., 1985), we examined effects of compounds ‘**8090** and ‘**8219** in the open field. We have reported previously that fluoxetine does not stimulate motor activities in WT and VMAT2 mice (Fukui et al., 2007). In the present study, the vehicle, 20 mg/kg fluoxetine, 2 mg/kg ‘**8090**, or 2 mg/kg ‘**8219** were administered (i.p.) to adult male and female C57BL/6J mice and their motor activities were followed over 36 min. Recall, in the tail suspension test mice were injected with fluoxetine or compounds and tested 30 min later over 6 min. An analysis of locomotor and rearing activities (Figure 7E-F) failed to detect any treatment effects over the first 30 min in the open field or in the subsequent 6 min. Importantly, neither fluoxetine nor either of the compounds exerted motor stimulating or inhibiting activities. Hence, the anti-depressive-like effects were not confounded by changes in motor responses. Taken together, our behavioral studies support anxiolytic and, more strongly, anti-depressant-like effects of these conformationally and sub-type selective SERT inhibitors.

## DISCUSSION

From large-library docking against the inward-open conformation of SERT emerged novel chemotypes with new pharmacology. In this still rare campaign against a transporter, the docking hit rate was high at 36%, as was the potency of the initial top-ranking molecules, many of which had sub-μM to even mid-nM Ki values. All of these compounds represent new chemotypes, topologically unrelated to known SERT inhibitors represented in the IUPHAR (Bröer, 2019; Southan et al., 2016) or ChEMBL (Gaulton *et al*., 2017) databases. The compounds were also selective for SERT (Figure S8), with little meaningful activity against well-known off-targets such as NET, DAT, and the serotonergic GPCRs and ion channels, in contrast with the broad promiscuity of ibogaine (Glick *et al*., 2001). This study thus supports the plausibility of targeting transporters for structure-based discovery. Several other observations merit emphasis. (**i**). The new chemotypes appear to stabilize a unique conformation of the transporter, closing the extracellular pathway without opening the intracellular pathway. How much this reflects the targeting of the inward-open state, against which the molecules were selected, and how much this reflects simply the novel chemotypes, is unclear, but it does present opportunities for new functional outcomes. (**ii**) Structure-based optimization improved potency, with the best molecule inhibiting SERT with Ki value of 3 nM. **(iii)** The cryo-EM structure of the SERT/**’8090**, while captured in the outward-open conformation owing to the conformational influence of Na^+^, broadly confirms the docked-predicted pose, with the new inhibitor binding site overlapping closely with the transporter’s central 5-HT binding site (Yang and Gouaux, 2021) and making predicted interactions, including with Asp98, Phe341, Phe335 and Y176 in both structures. (**iii**) The compounds were selected for favorable physical properties and have high brain exposure. This contributes to their potent activity in mouse anxiolytic and especially anti-depressant assays; in the latter, the new inhibitors were as much as 200-fold more potent than fluoxetine; this activity may reflect both their chemical novelty and the new states that they stabilize.

From a target perspective, the prioritization of potent initial actives against SERT suggests that docking will be fruitful against other members of this large and pharmacologically important family. Until now, most large library docking campaigns have targeted receptors such as GPCRs (Ballante *et al*., 2021; Kolb *et al*., 2009b; Roth *et al*., 2017; Sadybekov *et al*., 2020; Weiss *et al*., 2013), nuclear hormone receptors (Benod et al., 2013; Schapira et al., 2003), kinases (Bajusz et al., 2017; Muegge and Enyedy, 2004; Tavares et al., 2022) and soluble enzymes (Doman et al., 2002; Grüneberg et al., 2001; Lyu *et al*., 2019; Teotico et al., 2009); transporters have received relatively little attention. This study suggests that, with their well-formed binding sites, often evolved to accommodate metabolites and signaling molecules, they will make good targets for structure-based discovery. As with GPCRs and nuclear hormone receptors, the transporters adopt several key conformational states; these too may be individually targeted for different functional outcomes.

Differential phosphorylation of SERT conformations plays an important role in the regulation of transport activity (Zhang *et al*., 2016). A phosphorylation site (Thr276) sequestered from the cytoplasm in the outward-open conformation becomes accessible in the inward-open state (Coleman *et al*., 2019). Consequently, cocaine inhibits phosphorylation by stabilizing the outward-open state and ibogaine enhances phosphorylation by stabilizing the inward-open state (Zhang *et al*., 2016). In addition, SERT exists in a complex together with regulatory proteins including nNOS, PKG and PP2A, which may also be affected by SERT conformation (Chanrion et al., 2007; Zhang and Rudnick, 2011). Ligands that selectively open or close an individual pathway, such as **‘8090** and **‘8219**, may have selective effects, not only on SERT phosphorylation, but also on intracellular signaling mediated by these regulatory proteins. Conformation-selective ligands, by their selective action on SERT (directly) and associated proteins (indirectly) may have advantages over existing SERT inhibitors and may represent a more finely tuned approach to modifying SERT and associated signaling pathways.

Certain caveats should be mentioned. The large-scale docking was launched against the inward-open state conformation of SERT, seeking compounds that acted like ibogaine on SERT, but with higher target selectivity. Such compounds would allow testing the hypothesis that the unique conformational effect of ibogaine might be responsible for its reported ability to ameliorate opiate withdrawal (Alper *et al*., 1999; Brown, 2013; Schenberg *et al*., 2014; Wasko *et al*., 2018) and depression (Glick *et al*., 2001; Rodriguez *et al*., 2020). Our most potent ligands were not purely selective for this conformation, instead seeming to selectively stabilize a state in which the extracellular pathway is closed—as modeled—but the inward pathway is hardly changed. Moreover, the ligands differed markedly from ibogaine in their response to Na^+^ and Cl^-^ (Fig 3C or D). While the docking predicted structure interacts with the same residues and certainly the same overall site as that observed in the outward-*open* cryoEM structure, the superposition is only approximate, with an RMSD of 2.11 Å in ligand atoms (this improves to 1.17 Å when comparing the cryoEM structure to a docking to the outward-open conformation, Figure 5D). Even ibogaine, which strongly stabilizes inward-open outward-closed states, binds not only the inward-open state but also the outward-open and occluded states, reflecting overall similarities of the orthosteric sites in all three states (Coleman *et al*., 2019).

These cautions should not obscure the major observations from this study. Large library docking against the inward-open conformation of SERT discovered 13 new compounds representing 12 novel chemotypes that bound to the transporter with low μM to mid-nM concentrations. The docked structures templated optimization to the low nM range. As observed against other flexible receptors (Lyu *et al*., 2019; Manglik *et al*., 2016; Stein *et al*., 2020), the new chemotypes conferred new in vitro activities that likely contributed to the unusually high efficacy of the new SERT inhibitors in animal models of depression. Indeed, the selectivity of the new inhibitors for SERT versus off-targets like NET, DAT, and serotonergic GPCRs, and for a particular SERT conformational state, may make them useful as tool molecules to probe transporter function and therapeutic translation. Accordingly, we are making them openly available via the Millipore-Sigma probe collection. This study supports the pragmatism of structure-based campaigns against transporters and suggests that even for those as intensely studied as SERT, new structures can template the discovery of potent new chemotypes, conferring new pharmacology.

## METHODS

### Molecular Docking

Serotonin transporter bound to ibogaine in an inward-open conformation (Coleman *et al*., 2019) (PDB ID: 6DZZ) was used for docking a library of >200 million “lead-like” molecules from the ZINC20 database (http://zinc20.docking.org) using DOCK3.7 (Coleman et al., 2013). Forty five matching spheres or local hot-spots generated from the cryo-EM pose of ibogaine were used in the binding site for superimposing pre-generated flexible ligands and the poses were scored by summing the van der waals interaction energies, transporter-ligand electrostatics and ligand desolvation energies (Mysinger and Shoichet, 2010; Wei et al., 2002). The transporter atoms were protonated with Reduce (Word et al., 1999) and partial charges calculated using united-atom AMBER force field (D. A. Case et al., 2015). AMBER force field was also used for pre-generation of energy grids using QNIFFT (Gallagher and Sharp, 1998; Sharp, 1995) for Poisson–Boltzmann-based electrostatic potentials, CHEMGRID (Meng, 1992) for van der Waals potential, and SOLVMAP (Mysinger and Shoichet, 2010) for ligand desolvation.

The docking setup was optimized for its ability to enrich knows SERT binders including ibogaine, noribogaine, 5-hyroxytryptamine (5-HT), cocaine and known selective serotonin reuptake inhibitors (SSRIs) (Tatsumi *et al*., 1997), in favorable geometries with high complementarity versus a set of property matched decoys (Mysinger *et al*., 2012). About 50 decoys were generated for each ligand that had similar chemical properties to known ligands but were topologically dissimilar. The best optimized docking setup was evaluated using enrichment of ligands over decoys using log-adjusted area under the curve (logAUC values). The best docking setup was able to enrich the cryo-EM pose of ibogaine as well as dock other known ligands in the right conformation. All docked ligands were protonated with Marvin (version 15.11.23.0, ChemAxon, 2015; https://www.chemaxon.com) at pH 7.4, rendered into 3D with Corina (v.3.6.0026, Molecular Networks GmbH; https://www.mn-am.com/products/corina), and conformationally sampled using Omega (v.2.5.1.4, OpenEye Scientific Software;https://www.eyesopen.com/omega). Before launching a screen of >200 million make-on demand lead like molecules, an ‘extrema set’ of 61,687 molecules was docked in the optimized system to ensure that the molecules with correct physical properties were enriched.

Overall, in the prospective screen, each library molecule was sampled in about 4358 orientations, on average about 187 conformations were sampled over 5 days on 1000 cores. The top-ranking 300,000 molecules were filtered for novelty using ECFP4-based Tanimoto coefficient (Tc <0.35) against known inhibitors of SERT, DAT or NET and ∼28,000 annotated aminergic ligands acting at serotonin, dopamine and adrenergic receptors in ChEMBL (Gaulton *et al*., 2017). The remaining molecules were further clustered by an ECFP4-based Tc of 0.5. From the top 5,000 novel chemotypes, strained molecules with >2 kcal mol−1 internal strains were filtered out and the rest were visually inspected for the best docked poses with favorable interactions with the SERT active site, including salt bridge formation with Asp98, pi-pi stacking with Phe335 or Phe341 and polar interactions with Asn177. Ultimately, 49 molecules were selected for de novo synthesis and testing, out of which 36 were successfully synthesized and tested.

### Hit to lead optimization

Using 5 primary docking hits ‘9642, ‘6919, ‘2313, ‘4931 and ‘2305 as queries in SmallWorld (https://sw.docking.org/) and Arthor (http://arthor.docking.org) search engines (NextMove Software, Cambridge UK) (Irwin et al., 2020), substructure and similarity searches were conducted among >20 billion make-on-demand Enamine REAL molecules. The resulting analogs were further filtered based on Tc > 0.4 and docked to the SERT inward open active site. The docked poses were visually inspected for compatibility with the site, including salt bridge formation with Asp98, pi-pi stacking with Phe335 or Phe341 and polar interactions with Asn177 and prioritized analogs were synthesized and experimentally tested.

### Make-on-demand synthesis and Compound Handling

49 molecules from the large scale prospective docking were delivered within 5 weeks with a 73.4% fulfillment rate after a single synthesis attempt. These make-on demand molecules were derived from the Enamine REAL database (https://enamine.net/compound-collections/real-compounds). Portions of each compound (1 to 2 mg) were dissolved in DMSO at a concentration of 10 mM and stored as stock solutions at −20° C. The rest of the dry powder was also stored at −20° C for additional rounds of testing. Freeze-thaw cycles were minimized.

### Transport and efflux measurements

5-HT transport into RBL cells or SERT-transfected HeLa cells was measured as described previously (Zhang and Rudnick, 2005). Briefly, cells growing in 48- or 96-well plates were washed once with 100 µl (200 µl for 48-well plates) of PBS/CM (phosphate-buffered saline containing 0.1 mM CaCl2 and 1 mM MgCl2). 5-HT uptake assays were initiated by the addition of [^3^H]5-HT (20 nM final concentration) and unlabeled 5-HT to the indicated total concentration (for determination of K_M_ and Vmax). The assays were terminated after 10 min by three rapid washes with ice-cold PBS. The cells were then solubilized in 30 μl (150 μl for 48-well plates) of 0.01 M NaOH for 30 min. For 96-well assays, 120 µl of Optifluor (Perkin-Elmer) was added and accumulated [^3^H]5-HT was determined by liquid scintillation spectrometry in a PerkinElmer Microbeta plate counter. For 48-well assays, the NaOH-lysed cells were transferred to scintillation vials with 3 ml of Optifluor and counted by liquid scintillation spectrometry. For efflux experiments (in 48-well plates, incubations with 20 nM [^3^H]5-HT were extended to 15 min, followed by 2 washes with PBS/CM, and further incubation in 200 μl PBS/CM with or without the indicated inhibitor. Incubations were terminated as above at 5 min intervals up to 20 min and counted. Time courses were fitted by linear regression of the time courses.

### Binding measurements

Binding of the high-affinity cocaine analog [^125^I]β-CIT was measured in crude membrane preparations from SERT-transfected HeLa cells as described previously (Zhang and Rudnick, 2005). For membrane binding assays, frozen membranes from cells expressing SERT mutants were thawed on ice, applied to Multiscreen-FB 96-well filtration plates (Millipore, approximately 100 µg per well), and washed five times by filtration with 100 μl of binding buffer (10 mM HEPES buffer, pH 7.4, containing 150 mM NaCl, Na-isethionate, or NMDG-Cl as indicated). β-CIT binding was then initiated by the addition of 100 µl of binding buffer containing 0.1 nM [^125^I]β-CIT and the indicated concentration of individual test compounds. Binding was allowed to proceed for 1.5 h at 20°C with gentle rocking. The reaction was stopped by filtration and three washes with 100 μl of ice-cold PBS buffer. 50 μl of Optifluor was added to each filter and the plates were counted with a PerkinElmer Microbeta plate counter.

### Accessibility measurements

Conformational changes were measured using the accessibility of cysteine residues placed in the cytoplasmic (S277C) and extracellular (Y107C) permeation pathways as described (Jacobs *et al*., 2007a). For measuring extracellular pathway accessibility, intact cells expressing SERT C109A-Y107C growing in 96-well plates were incubated for 15 min with 0.01-10 mM MTSET in the presence or absence of test compounds and then washed with PBS/CM. Transport rates were then measured to determine the reactivity of Y107C under the experimental conditions tested. For cytoplasmic pathway accessibility, membranes prepared from cells expressing SERT S277C-X5C were applied to Multiscreen-FB 96-well filtration plates, washed as in the binding assays, and then incubated for 15 min with 0.01-1 mM MTSEA in the presence or absence of test compounds. and then washed with PBS/CM. Binding of the high affinity cocaine analog β-CIT was measured on the filters as described above. Modification of Cys277 inactivates β-CIT binding by preventing closure of the cytoplasmic pathway and opening of the extracellular pathway (Zhang and Rudnick, 2006). For measurements with both transport and binding, the concentration of MTSEA or MTSET leading to half-maximal inactivation were used to calculate the rate constant for inactivation and expressed either as that rate constant or the rate relative to the control rate in the absence of test compound. All test compounds and other inhibitors were added at 10x the K_I_ measured for inhibition of binding or transport, depending on the assay.

### Dissociation measurements

Dissociation rates for **‘8090** and **‘8219** were determined under whole-cell patch clamp by first inhibiting SERT with a saturating concentration of the inhibitor and then washing the inhibitor away while testing for recovery of SERT currents by frequent application of 5-HT. HEK293 monoclonal cells stably expressing GFP-hSERT were incubated for 2h with 75nM and 50nM of **‘8090** and **‘8219** respectively in an external solution (140mM NaCl, 3mMKCl, 2.5mM CaCl2, 2.0mM MgCl2, 10mM HEPES pH 7.4, 20mM glucose). Patch-clamp pipettes were back-filled with an internal solution (133 mM K-MES, 1.0mM CaCl_2_, 0.7mM MgCl_2_, 10mM EGTA, 10mM HEPES pH 7.2). Patched cells were constantly perfused for 37.5min with an inhibitor-free external solution. A 1s pulse of 10μM 5-HT was applied to the cell every 15s during this perfusion. The amplitude of 5-HT induced steady-state current (Schicker et al., 2012) was recorded and plotted vs. time.

### Expression and purification of SERT and Fab15B8

The ΔN72/ΔC13 N- and C-terminally truncated human wild-type SERT amino acid sequence (Coleman et al. 2019) was codon optimized and synthesized (Twist Bioscience). The SERT gene (*Slc6a4*) fragment was cloned into a pcDNA3.4-zeocin-TetO vector with a C-terminal human rhinovirus 3C cleavage site, followed by a human protein C tag (EDQVDPRLIDGK), and a 10 x polyhistidine tag. Expi293F cells at 3.0 x 10^6^ cells/ml were transfected with 1 µg DNA per ml culture using the Expi293 Expifectamine kit (Life Technologies) according to the manufacturer’s recommendations. The following day, expression was induced with 1 µg/ml doxycycline hyclate (Sigma Aldrich), and Expifectamine Enhancer Solutions I and II (Life Technologies) were added. After 48 h cells were harvested by centrifugation at 4,000 x *g*, and stored at −80°C until further use. On the day of purification, cells were thawed and dounce homogenized into ice cold solubilization buffer comprised of 100 mM NaCl, 20 mM HEPES pH 7.50, 1% (w/v) DDM, 0.1% (w/v) CHS, EDTA-free protease inhibitor (Thermo Fisher), and 30 µM compound **‘8090**. Cells were solubilized for 60 min at 4°C and centrifuged at 14,000 x g for 30 min at 4 C. The supernatant was supplemented with 2 mM CaCl_2_ and loaded over homemade anti-protein C affinity resin. The resin was washed with 25 C.V. of buffer comprised of 100 mM NaCl, 20 mM HEPES pH 7.50, 0.1% (w/v) DDM 0.01% (w/v) CHS, 1 mM CaCl_2_, and 10 µM **‘8090**. SERT was eluted with 100 mM NaCl, 20 mM HEPES pH 7.50, 0.1% (w/v) DDM 0.01% (w/v) CHS, 1 mM EDTA, 0.2 mg/ml protein C peptide (Genscript), and 30 µM **‘8090**.

Fab15B8 heavy and light chain amino acid sequences were obtained from the literature (Coleman *et al*., 2019) and codon optimized genes were designed with an N-terminal H7 signal sequence and a C-terminal polyhistidine tag. The genes were synthesized and cloned into the commercially available pTWIST-CMV expression vector (Twist Bioscience). Expi293F cells at 3.0 x 10^6^ cells/ml were transfected with 1 µg total DNA per ml culture of heavy chain and Light chain DNA in a 2:1 mass ratio, using the Expi293 Expifectamine kit (Life Technologies). The following day, Expifectamine Enhancer Solutions I and II (Life Technologies) were added. At approximately 120 h, culture was harvested by centrifugation at 4,000 x *g* and the supernatant containing Fab15B8 was loaded over a Ni-NTA column. The column was extensively washed with 20 mM HEPES pH 7.50, 100 mM NaCl, 30 mM imidazole. Fab15B8 was eluted with 20 mM HEPES pH 7.50, 100 mM NaCl, 250 mM imidazole, concentrated on a 10k MWCO spin filter (Amicon), and aliquots were flash frozen in liquid N_2_ and stored at −80°C until use.

### Nanodisc-SERT reconstitution

Approximately 150 µg of purified SERT was reconstituted into lipidic nanodiscs by mixing with purified MSPNW11 and a lipid mixture containing 2:3 weight ratio of 1-palmitoyl-2-oleoylphosphatidylcholine (POPC, Avanti) and 1-palmitoyl-2-oleoyl-*sn*-glycero-3-phospho-(1′-rac-glycerol) (POPG, Avanti). A SERT:MSPNW11:lipid molar ratio of 1:20:800 was used in buffer comprising 20 mM HEPES pH 7.50, 100 mM NaCl, 1 mM EDTA, 30 µM **‘8090**. MSPNW11 was purified as described previously (Billesbølle et al., 2020). Samples were incubated under slow rotation for 1 h at 4°C. Detergent was removed by addition of 200 mg/ml SM-2 BioBeads (BioRad) followed by incubation under slow rotation for 16 h at 4°C. The reconstituted sample was separated from BioBeads and 2 mM CaCl_2_ was added before loading over an anti-protein C affinity resin. The resin was washed with 100 mM NaCl, 20 mM HEPES (pH 7.50), 1 mM CaCl_2_ and 10 µM **‘8090**. Nanodisc-SERT was eluted with 100 mM NaCl, 20 mM HEPES pH 7.50, 0.5 mM EDTA, 0.2 mg/ml protein C peptide (Genscript), and 30 µM **‘8090**. The nanodisc-SERT sample was concentrated using a 50k MWCO spin filter (Amicon).

### Cryo-EM sample preparation and data collection

Nanodisc reconstituted SERT was mixed with 1.25 fold molar excess purified Fab15B8 and incubated 30 min on ice. The complex was purified by size exclusion chromatography (SEC) on a Superdex S200 Increase 10/300 GL column (GE Healthcare) into buffer comprised of 100 mM NaCl, 20 mM HEPES pH 7.50, and 10 µM **‘8090**. Peak fractions containing the monomeric SERT complex were supplemented with **‘8090** to 30 µM and concentrated to ∼10 µM with a 50k MWCO spin filter (Amicon). Monodispersity of the final EM sample was assessed by analytical fluorescence SEC using ∼ 5 µg protein and the chromatography buffer and column. Tryptophan fluorescence was recorded with an FP-1520 Intelligent Fluorescence Detector (Jasco) using λ_ex_ = 280 nm and λ_em_ = 350 nm.

The 2.5 µl sample was applied to glow discharged UltrAuFoil (R 1.2/1.3) 300 mesh grids (Qauntifoil). Grids were plunge vitrified into liquid ethane using a Vitrobot Mark IV (Thermo Fisher) with 5 s wait time, 3-5 s blot time, and 0 blot force. The blotting chamber was maintained at 100% humidity and 22°C. Vitrified grids were clipped with Autogrid sample carrier assemblies (Thermo Fisher) immediately prior to imaging.

Movies of **‘8090**-bound MSPNW11-SERT-Fab15B8 embedded in ice were recorded using a Titan Krios Gi3 (Thermo Fisher) equipped with a BioQuantum Energy Filter (Gatan) and a K3 Direct Electron Detector (Gatan). Data were collected using Serial EM (Mastronarde, 2003) running a 3 x 3 image shift script at 0° stage tilt. A 105,000 x nominal magnification with 100 µm objective aperture was used in superresolution mode with a physical pixel size of 0.81 Å pixel^-1^. Movies were recorded using dose fractionated illumination conditions with a total exposure of 50.0 e^-^ Å^-2^ delivered over 60 frames yielding 0.833 e^-^ Å^-2^ frame^-1^.

### Data processing

Raw movies were imported into cryoSPARC v3.2.0 (Structura Biotechnology), patch motion corrected with 0.5 Fourier cropping, and contrast transfer functions were calculated for the resulting micrographs using Patch CTF Estimation. Particles were template picked using an ab initio model that was generated during data collection in cryoSPARC Live (Structura Biotechnology). A total of 3,313,742 particles were extracted with a 360 pixel box that was binned to 96 pixels, and sorted by two rounds of 3D classification by heterogenous refinement, using initial model templates low pass filtered to 20 Å. Particles were unbinned and subjected to an additional round of sorting by heterogenous refinement, using two initial models of SERT. Finally non-uniform refinement was performed, before particles were exported using the pyem v0.5 (Asarnow, 2019). An inclusion mask covering SERT was generated with the Segger tool in Chimera and the mask.py tool in pyem v0.5. Particles and mask were imported into Relion v3.0 and subjected to 3D classification without image alignment. A series of classifications were performed varying the number of classes and the T factor. The resulting 187,696 particles were brought back into cryoSPARC and non-uniform refinement followed by local refinement using a mask covering SERT and the variable chains of Fab15B8.

A Directional Fourier shell correlation (dFSC) was calculated using half maps and the final output mask from the local refinement (Dang et al., 2017). Local resolution estimation were calculated in cryoSPARC. Euler angle distribution was visualized with the star2bild.py script. Model building and refinement were carried out using PDB 6VRH as starting model, which was fit into the 3.0 Å SERT map using ChimeraX (Goddard et al., 2018). A rough model was generated using ISOLDE extension (Croll, 2018) which was further refined by iterations of real space refinement in Phenix (Adams *et al*., 2010) and manual refinement in Coot (Emsley and Cowtan, 2004). The **‘8090** model and rotamer library were generated with PRODRG server (Schüttelkopf and van Aalten, 2004), and docked using Coot. Final map-model validations were carried out using Molprobity and EMRinger in Phenix.

### Animals

Animal experiments were conducted under protocols approved by the UCSF Institutional Animal Care and Use Committee (IACUC) and by the Duke University IACUC and were conducted in accordance with the NIH Guide for the Care and Use of Laboratory animals. Adult (8-10 weeks old) male C56BL/6J mice were purchased from the Jackson Laboratory (strain #000664). The experiments testing antidepressant drugs/compounds were conducted with adult (4-6 mos) male and female wild-type and vesicular monoamine transporter 2 heterozygous mice. The open field studies were conducted with adult (4 mos) male and female C57BL/6J mice. All mice were housed 3-4 in cages, in a humidity and temperature-controlled room, on a 12:12 h light/dark cycle (UCSF) or on a 14:10 h light/dark cycle (lights on 0600 h) with food and water provided *ad libitum*. The antidepressant and open field experiments were conducted between 1000 and 1500 h.

### Drugs and Compounds

All novel ligands were synthesized by Enamine to ≥95% analytic purity. **‘8219** was resuspended in 20% cyclodextran and **‘8090** in NaCl 0.9%. in the anxiety assays. In the tail suspension and open field tests, the vehicle consisted of N,N-dimethylacetamide (final volume 0.5%; Sigma-Aldrich) brought to volume with 5% 2-hydroxypropoyl-β-cyclodextrin (Sigma-Aldrich) in water (Mediatech Inc., Manassas, VA). As a positive control in tail suspension, fluoxetine (Sigma-Aldrich) was used.

### Elevated Plus-Maze Assay

The plus-maze apparatus consisted of two open arms and two closed arms (with 15 cm high, opaque walls) of the same dimensions (35 x 9 cm), elevated to a height of 50 cm. Mice received 100 μl of each compound (10 mg/kg; i.p.) or vehicle (saline for **‘8219** and 20% cyclodextran for **‘8090**) once a day for 10 consecutive days. After the last injection (on the 10^th^ day), mice were placed in a Plexiglas cylinder for 30 minutes before being placed into the center area of the plus-maze, facing a closed arm. A camera placed ∼ 2 m above the maze recorded the amount and the number of times each mouse entered each arm, over a period of 5 min. The total distance travelled in the apparatus was also recorded. A single injection of paroxetine (10 mg/kg. i.p.) was used as positive control. Animals receiving paroxetine were also first habituated for 30 minutes in a Plexiglas cylinder before entering the plus-maze. All statistical analyses were performed with Prism (Graph Pad). Data are reported as mean +/- SEM. In all experiments, unpaired Student’s t-test was used to compare the effect of the compounds and paroxetine with their vehicle control.

### Tail Suspension, Open Field Tests, and Statistics

This test was conducted using the Med Associates (ST. Albans, VT) tail suspension apparatus where the body weight of the mouse was used as a control for the force of struggle activity as described (Fukui *et al*., 2007). All drugs or compounds were injected (i.p.) in a 5 mL/kg volume. Wild-type (WT) and VMAT2 heterozygous (HET) mice were administered the vehicle, 20 mg/kg fluoxetine (Sigma-Aldrich), and different doses of 2237 or 8507. The mice were tested acutely 30 min and 24 h later in the tail suspension assay over 6 min for each day.

Locomotor activities were evaluated acutely in an open field (21 x 21 x 30 cm; Omnitech Electronics, Columbus, OH) illuminated at 180 lux (Fukui *et al*., 2007) with adult male and female C57BL/6J. Mice were injected (i.p.) with the vehicle, 20 mg/kg fluoxetine (Sigma-Aldrich), 2 mg/kg 2237, or 2 mg/kg 8507 (all in a 5 mL/kg volume) and placed immediately into the open field. Locomotion (distance traveled) was monitored over 30 min using Fusion Integra software (Omnitech) in 5-min blocks or as cumulative activities.

The data are presented as means ± standard errors of the mean and were analyzed by one-way ANOVA or repeated measures ANOVA followed with Dunnett or Bonferroni corrected pair-wise comparisons (IBM SPSS Statistics 27 programs; IBM, Chicago, IL). A p<0.05 was considered statistically significant.

### Neurotransmitter Transporter Assays

DAT, NET, and SERT activities were determined using the neurotransmitter transporter update assay kit from Molecular Devices (Category R8174). Briefly, HEK293 cells stably expressing human DAT, NET, or SERT were plated in Poly-L-Lys (PLL) coated 384-well black clear bottom plates in DMEM supplemented with 1% dialyzed FBS (dFBS), at a density of 15,000 cells in 40 µl per well. After overnight recovery, the cells were removed of medium, received 25 µl per well drug solutions prepared in assay buffer (1x HBSS, 20 mM HEPES, pH 7.40, supplemented with 1 mg/ml BSA) for 30 min at 37°C, followed by 25 µl per well of dye solution for an additional 30 min incubation at 37°C. Cocaine, Nisoxetine, and fluoxetine served as positive controls for DAT, NET, and SERT, respectively. Fluorescence intensity was measured on the FlexStation II with excitation at 440 nm and emission at 520 nm. Relative fluorescence units (RLU) were exported and analyzed in Prism 9.0.

### GPCRome screening assays

Off-target agonist activity at human GPCRome was carried out using the PRESTO-Tango assays as described before (Kroeze *et al*., 2015) with modifications. In detail, HTLA cells were plated in PLL coated white clear-bottom 384-well plates in DMEM supplemented with 1% dFBS, at a density of 10,000 cells in 40 µl per well. After recovery for about 6-hour, the cells were transfected with 20 ng DNA per well for overnight incubation, followed by addition of 10 µl of selected test compounds, prepared in DMEM supplemented with 1% dFBS. The cells were incubated with drug overnight (usually 16 – 20 hours). Medium and drugs were then removed and 20 µl per well of BrightGlo reagents diluted in assay buffer were added. Plates were incubated for 20 min at room temperature in the dark and luminescence was counted. In each plate, the dopamine receptor D_2_ was included as an assay control and was stimulated with 100 nM of the agonist quinpirole. Each receptor had 4 replicate wells of basal (with medium) and 4 replicate wells of sample (10 µM final in this case). Results were expressed in as fold change over average basal.

### GPCR β-Arrestin Tango Assays

HTLA cells were transfected with target receptor construct in DMEM supplemented with 10% FBS for overnight, plated in PLL coated white clear-bottom 384-well plates in DMEM supplemental at a density of 10,000 cells in 40 µl per well. After recovery for about 6-hours, compound in serial dilutions, prepared in DMEM supplemented with 1% dFBS, were added to cells, 10 µl per well in 5x of the final designed. The plate was incubated overnight, usually 16 – 20-hours and luminescence counts were determined as above. Results were analyzed and processed in Prism 9.0.

### GloSensor cAMP Assays

HEK293 T cells were co-transfected with the target receptor construct and GloSensor cAMP reporter (Promega) in DMEM supplemented with 10% FBS for overnight incubation. Transfected cells were plated in PLL coated white clear-bottom 384-well plates in DMEM supplemented with 1% dFBS at a density of 15,000 to 20,000 cells in 40 µl per well. After recovery of a minimum of 6-hours (up to 24-hours), cells were used for GloSensor cAMP assays. Briefly, cells were removed of medium and received 25 µl per well compound solutions, prepared in assay buffer (1x HBSS, 20 mM HEPES, pH 7.40, supplemented with 1 mg/ml BSA) supplemented with 4 mM luciferin. For G_s_ agonist activity, the plates were counted for luminescence after 20 min incubation at room temperature in the dark. For G_i_ agonist activity, 10 µl of isoproterenol (ISO at a final of 100 nM, used to activate endogenous ß_2_-receptors to activate G_s_ and then adenylyl cyclase activity) was added at 15 min after compound and the plate was counted after 20 min as above. Results were analyzed in Prism 9.0.

