## Supplementary Figures and Tables for "Structure-based Discovery of Conformationally Selective Inhibitors of the Serotonin Transporter"

**Contents**:

**Figure S1.** Structure based strategy for discovery of SERT inhibitors with novel chemotypes, related to Figure 1 and STAR Methods.

**Figure S2.** Analogs of top five hits, related to Figures 1 and 2.

**Figure S3.** Docked poses of **'8219** (R-enantiomer) and **'8221** (S-enantiomer), related to Figure 2.

**Figure S4**. Inhibition of Transport and binding by **'8090** and **'8219** and kinetics of dissociation, measured by recovery of 5-HT-dependent current during washout of **'8219** and **'8090.** related to Figure 3 and STAR Methods.

**Figure S5.** Dependence of **'8090** binding on $Na^+$ and $Cl^-$, related to Figure 4 and STAR Methods.

**Figure S6**. Representation of SERT in two conformations with reactive cysteines shown, related to Figure 4.

**Figure S7.** Cryo-EM analysis of **'8090** bound SERT, related to Figure 5.

**Figure S8.** Comprehensive activity profiling against NET, DAT and SERT and GPCRome, related to Figure 2 and STAR methods.

**Figure S9.** Pharmacokinetic analysis of **'8090** and **'8219**, related to Figures 6 and 7 and STAR methods.

**Table S1**. Active docking hits that inhibited greater than 50% of [$^3$H]5-HT transport by SERT, related to Figure 1, S1 and STAR methods.

**Table S2**. Active analogs of top five hits tested at SERT, related to Figure 1, S1 and STAR methods.

**Table S3**. Cryo-EM data collection, refinement and validation statistics, related to Figure 5 and Figure S7.

**Table S4**. Affinity differs between binding and transport assays, related to Figure 3 and STAR methods.

**Table S5**. Calculated binding and dissociation rates for '8219 and '8090, related to Figure 3 and STAR methods.

**Table S6.** Dependence of affinity on $Na^+$ and $Cl^-$ ions, related to Figure 3 and STAR methods.

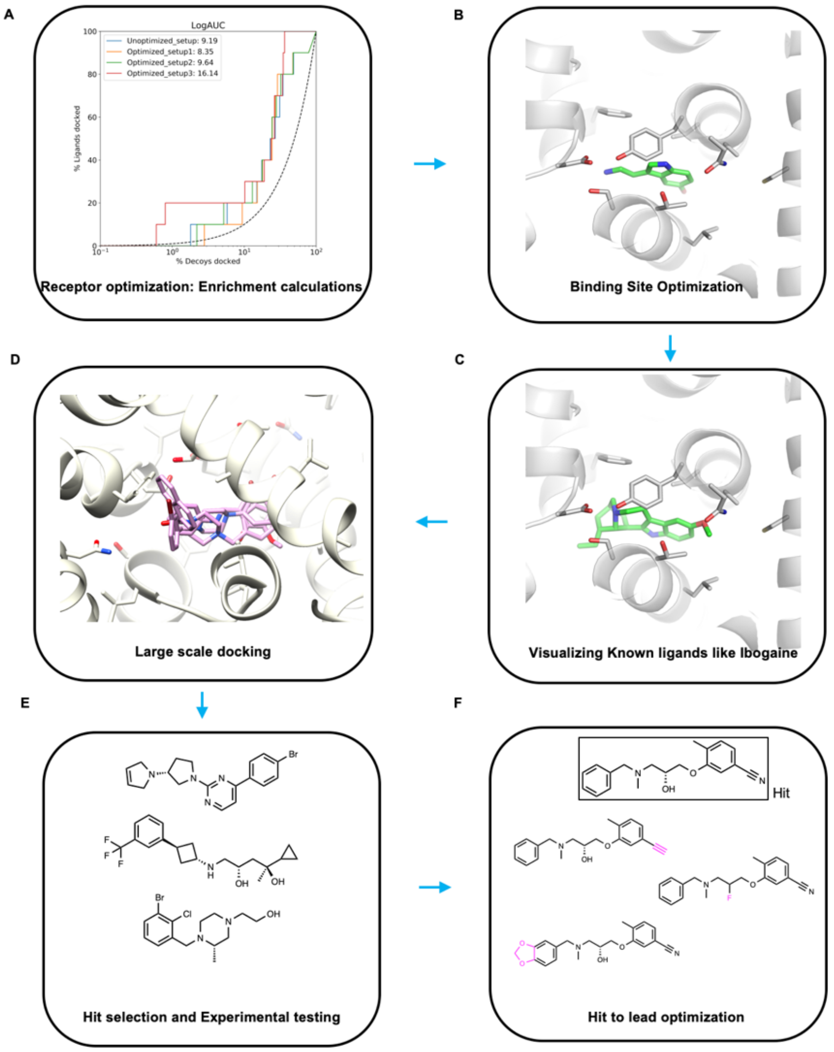

**Figure S1. Structure based strategy for discovery of SERT inhibitors with novel chemotypes, related to Figure 1 and STAR Methods**

(**A**) The inward open state conformation of SERT was evaluated for its ability to enrich known SERT ligands over property-matched decoys through docking to the orthosteric site, using DOCK 3.7. The system was optimized using addition of dielectric and ligand desolvation terms to get better early enrichments. (**B**) The binding site was further optimized using energy minimization. The optimized transporter was further analyzed by re-docking both ligands and decoys using DOCK 3.7 and calculating enrichments. (**C**) The binding poses of known ligands were visualized for optimal binding in the orthosteric pocket. (**D**) Over 200 million "lead-like" molecules from ZINC20 were prospectively screened against the inward open state of the transporter by molecular docking. (**E**) The top 300,000 were further filtered for novelty against all known SERT binders (Tc < 0.30) as well as ~28000 annotated aminergic ligands. Top 10,000 molecules were further visualized for favorable geometries and interactions with key binding residues including salt bridge formation with Asp98 and stacking with Phe 341. 36 compounds were tested in inhibition assays. (**F**) The top five active compounds were optimized for potency and selectivity against the inward-open state through both docking and experimental testing.

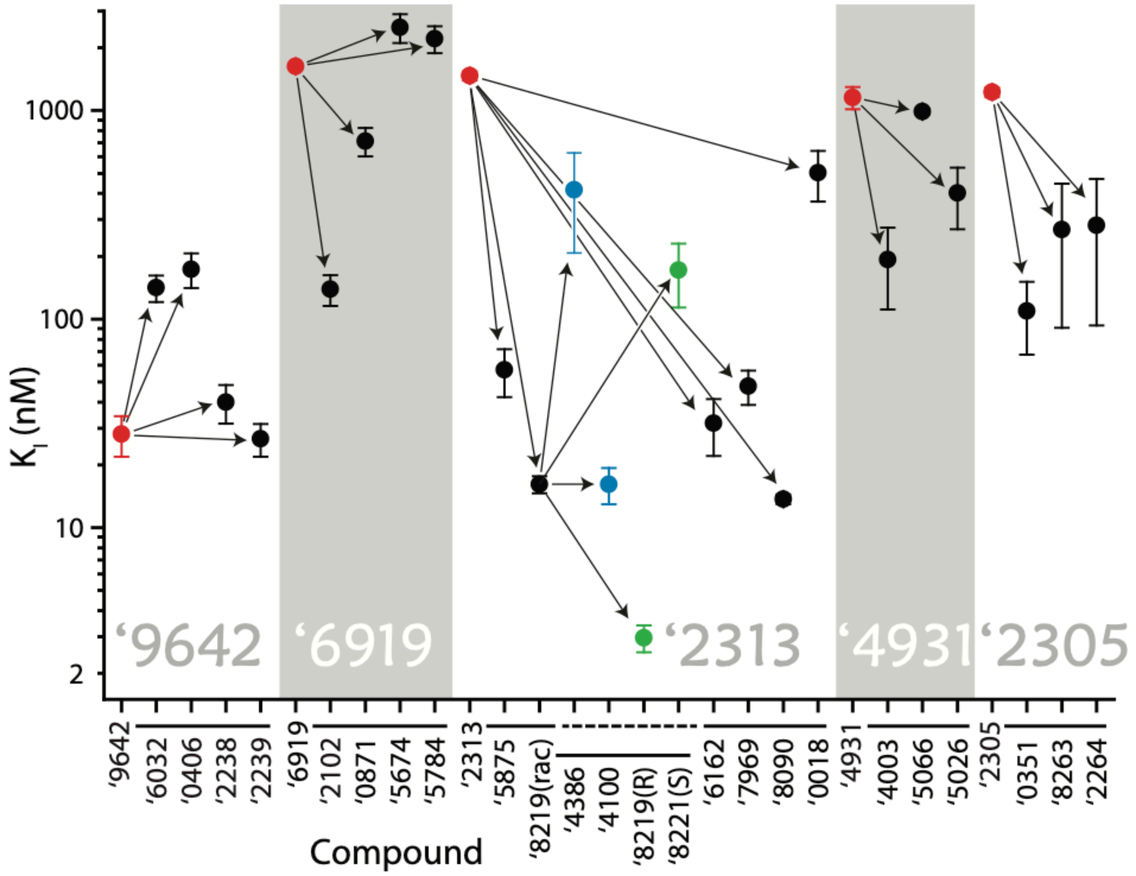

**Figure S2. Analogs of top five hits, related to Figures 1 and 2.**
Red symbols represent initial hits (Table S1) Black symbols represent analogs of the initial hits, Blue symbols represent analogs of **‘8219**, and green symbols represent the individual (R) and (S) isomers, **‘8219** and **‘8221**, respectively. **‘8219** (rac) is the racemic mixture of these isomers (Table S2).

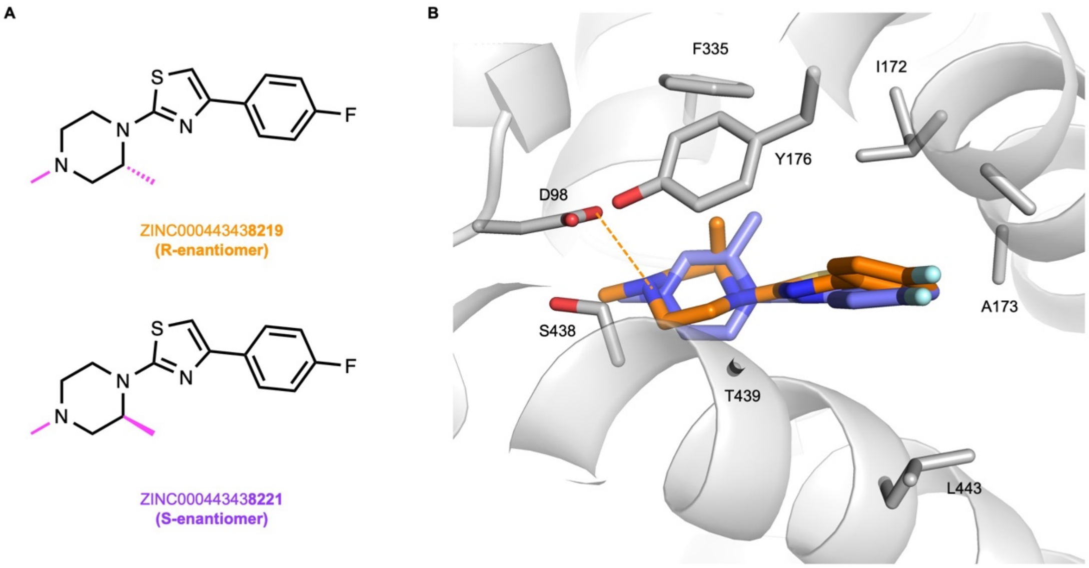

**Figure S3. Docked poses of '8219 (R-enantiomer) and '8221 (S-enantiomer), related to Figure 2.**

**(A)** Chemical structures of the two enantiomers of the same compound. (B) Docked poses of the **'8219** (R-enantiomer) and **'8221** (S-enantiomer) with carbons represented in orange and blue respectively. **'8221** is unable to engage the key residue D98 due to oreintation of its methyl group.

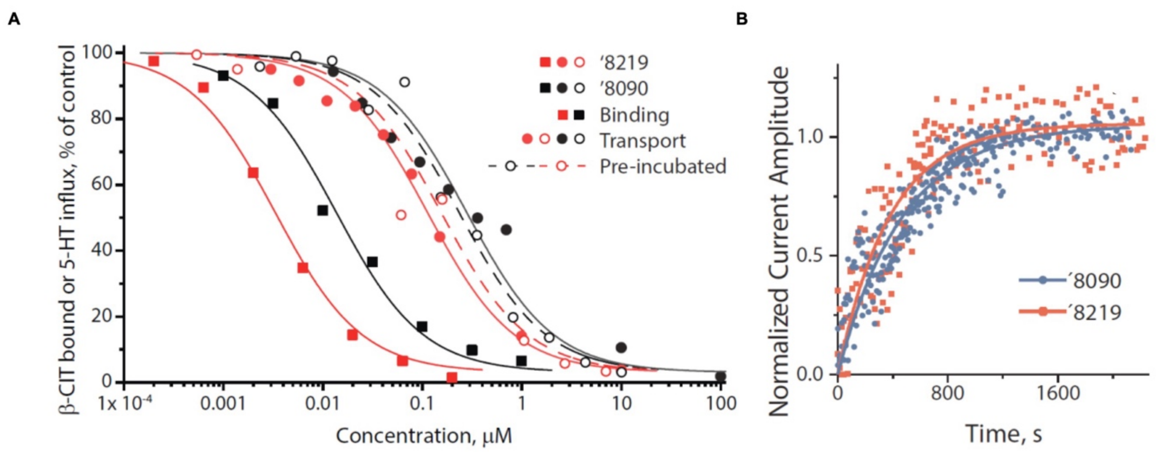

**Figure S4. Inhibition of Transport and binding by '8090 and '8219 and kinetics of dissociation, related to Figure 3 and STAR Methods.**
**(A)** Percentage of remaining [$^{125}$I]β-CIT binding (squares) or [$^{3}$H]5-HT transport (circles) as a function of **'8219** (red symbols) or **'8090** (black symbols) concentration. [$^{3}$H]5-HT was added simultaneously with Inhibitors (solid lines, filled circles) or after a 30 min preincubation (dashed lines, open circles). **(B)** Time course for recovery of 5-HT-induced currents. HEK-293 cells expressing hSERT were pre-incubated for 60 m with 50 nM **'8219** or 75 nM **'8090** patched, and superfused with inhibitor-free buffer. At intervals of 15 s, the cell was exposed to a brief pulse of 5-HT and the increase in inward current amplitude measured. The compounds dissociated at rates of $2.27 \pm 0.20 \times 10^{-3} s^{-1}$ and $1.82 \pm 0.09 \times 10^{-3} sec^{-1}$ for **'8219** and **'8090**, respectively. Using $K_D$ values of 5 nM for **'8219** and 15 nM for **'8090**, we calculated that $k_{on}$ rates would be 0.045 and 0.012 $M^{-1}$ $s^{-1}$ for **'8219** and **'8090**, respectively. At concentrations giving half-maximal inhibition, the half time for equilibration of binding should be less than 7 minutes for both compounds (Table S5), much shorter than the 30 min pre-incubation with inhibitor, which did not lead to a measureable difference in $IC_{50}$ values relative to adding substrate and inhibitor simultaneously in the transport experiments.

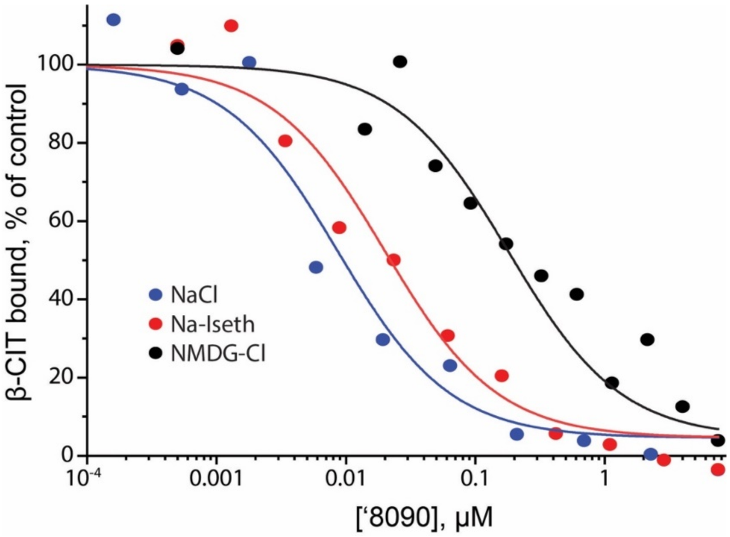

**Figure S5. Dependence of '8090 binding on $Na^+$ and $Cl^-$, related to Figure 4 and STAR Methods.**

$Na^+$ increased **'8090** affinity in equilibrium displacement of $[^{125}I]$β-CIT. Membranes from cells expressing SERT were incubated with ? nM $[^{125}I]$β-CIT and the indicated concentrations of **'8090** in PBS (control), PBS in which $Na^+$ was replaced with $NMDG^+$ (black line and circles) or $Cl^-$ was replaced with isethionate (red line and circles). The presence of $Cl^-$ increased **'8090** inhibitory potency less than 2-fold (not significant), from a $K_I$ of 23 ± 11 nM to 16 ± 0.4 nM in these experiments (n-3). $Na^+$ increased **'8090** inhibitory potency 11-fold, from a $K_I$ of 176 ± 46 nM to 16 ± 6 nM.

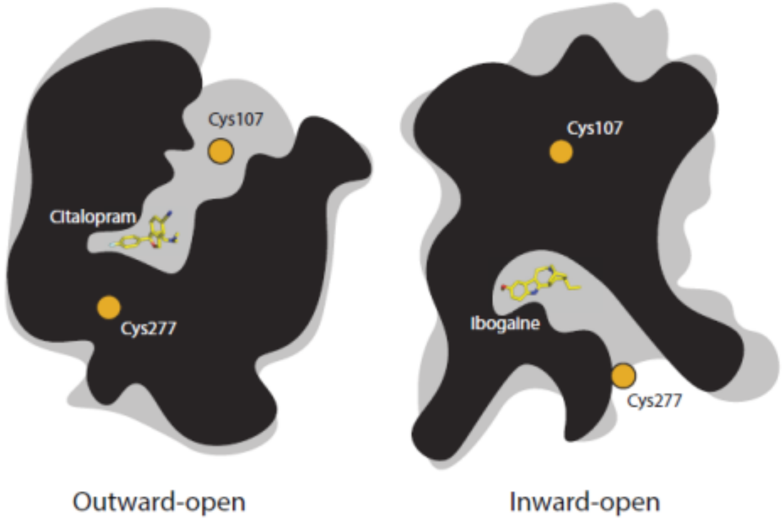

**Figure S6**. **Representation of SERT in two conformations with reactive cysteines shown, related to Figure 4.**
The outward-open conformation is shown with Cys107 exposed in the extracellular pathway, residue 277 buried and citalopram at the central substrate site (PDB ID: 5I73) (Coleman et al., 2016). The inward-open conformation is shown with Cys277 exposed in the cytoplasmic pathway, residue 107 buried and ibogaine in the central substrate (PDB ID: 6DZZ) (Coleman et al., 2019).

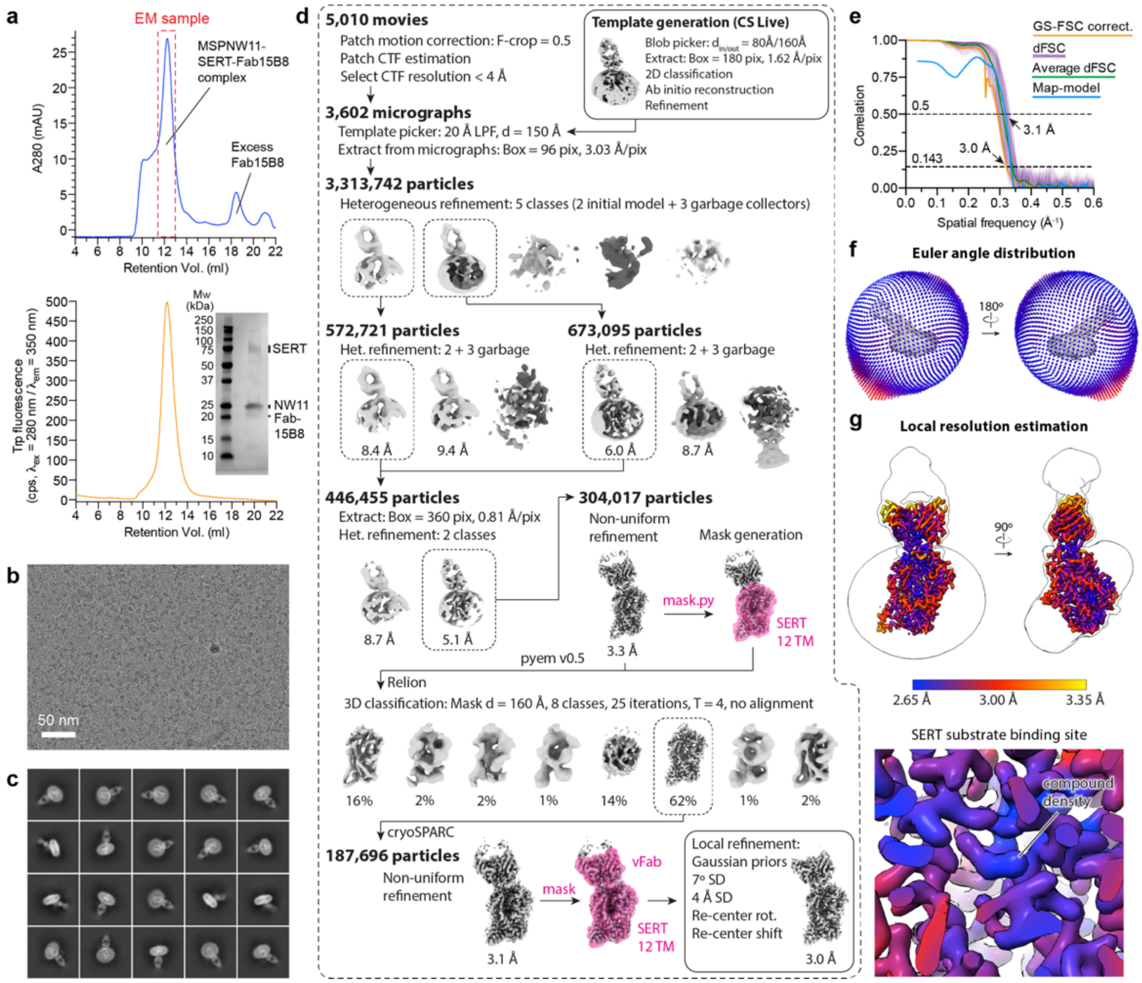

**Figure S7. Cryo-EM analysis of ‘8090 bound SERT, related to Figure 5**
**(A)** Size exclusion chromatogram showing purification of nanodisc-reconstituted SERT in complex with Fab15B8 and compound **‘8090** in the presence of $Na^+$. The final concentrated sample was analyzed by tryptophan fluorescence SEC and SDS-PAGE prior to vitrification. **(B)** Representative good motion-corrected micrograph showing monodisperse nanodisc-SERT-Fab15B8 particles. **(C)** Select 2D class averages generated from the fully processed dataset showing distinct views of nanodisc-SERT-Fab15B8. **(D)** The empirically determined image processing workflow for nanodisc-SERT-Fab15B8. Processing was achieved by combining tools in cryoSPARC v3.2.0 and Relion v3.0, enabled by the pyem v0.5 script package (Asarnow, 2019). Particle numbers for each step are provided and resolutions are FSC = 0.143 cut-off values reported in cryoSPARC. **(E)** Directional Fourier shell correlation (dFSC) curves from the final nanodisc-SERT-Fab15B8 reconstruction. Map-model FSC curves calculated in Phenix (Adams et al., 2010.)**(F)** Euler angle distribution from the final reconstruction visualized as two orthogonal views using the star2bild.py script. **(G)** Local resolution estimation calculated in cryoSPARC, shown in the same orthogonal views as f. A detailed view of the SERT substrate binding site is also shown.

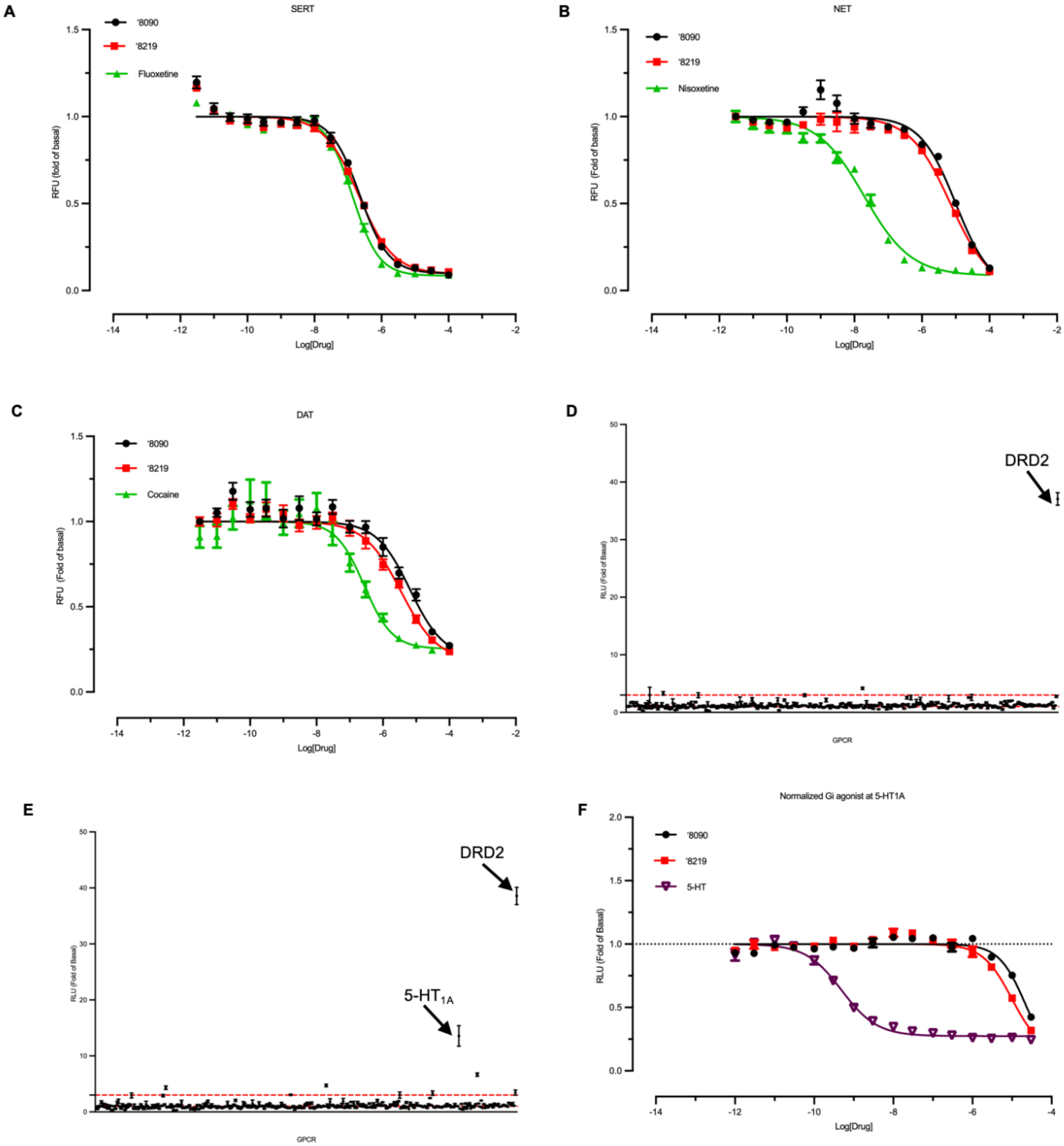

**Figure S8. Comprehensive activity profiling against NET, DAT and SERT and GPCRome, related to Figure 2 and STAR methods.**
(**A-C**) Measurement of inhibitory activity of **'8090** and **'8219** at SERT (**A**), NET (**B**) and DAT (**C**) using cell based functional assay. (**D-E**) Tango β-arrestin 2 recruitment assay testing the activity of **'8219** (**D**) and **'8090** (**E**) on >300 GPCRs to assess off-target activity. Both compounds were tested at 10µM with dopamine D2 receptor (DRD2) as control.
(**F**) Confirmation of primary screen hit for **'8090** as concentration dose response curve.

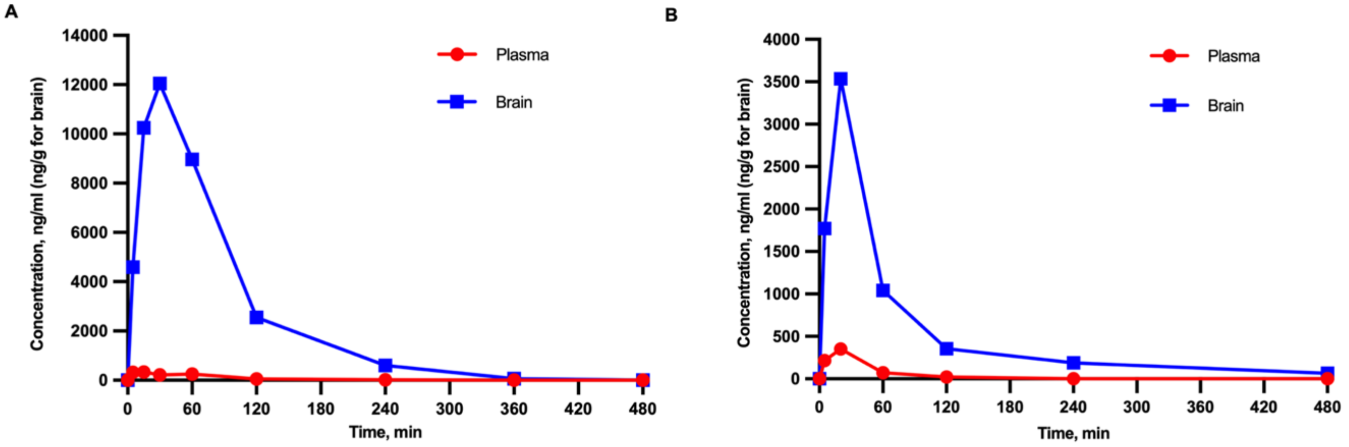

**Compound ‘8090, 10 mg/kg dose, IP administration**

| Pharmacokinetic Parameters | Brain | Plasma |
|---|---|---|
| $T_{1/2}$ | 44.3 min | 47.2 min |
| $C_{max}$ | 12000 ng/g | 332 ng/ml |
| AUC | 1150000 ng*min/g | 29000 ng*min/ml |

**Compound ‘8219, 10 mg/kg dose, IP administration**

| Pharmacokinetic Parameters | Brain | Plasma |
|---|---|---|
| $T_{1/2}$ | 147 min | 25.9 min |
| $C_{max}$ | 3530 ng/g | 352 ng/ml |
| AUC | 241000 ng*min/g | 16900 ng*min/ml |

**Figure S9. Pharmacokinetic analysis of ‘8090 and ‘8219, related to Figures 6 and 7 and STAR methods.**
(**A**) Concentration-time curve for **‘8090** in mice following IP dosing at 10 mg/kg.
(**B**) Concentration-time curve for **‘8219** in mice following IP dosing at 10 mg/kg.

**Table S1. Active docking hits that inhibited greater than 50% of [$^3$H]5-HT transport by SERT, related to Figure 1, S1 and STAR methods.**

| ZINC-ID | Chemical Structure | Ki, µM | SD |
|---|---|---|---|
| ZINC000623997601 | 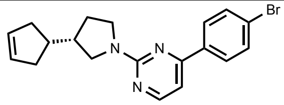 | 12.5 | 2.5 |
| ZINC000305339642 | 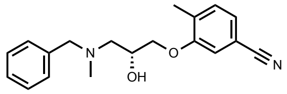 | 0.029 | 0.010 |
| ZINC000853098204 | 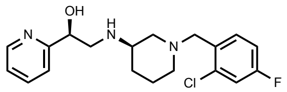 | 3 | 1 |
| ZINC000623756919 | 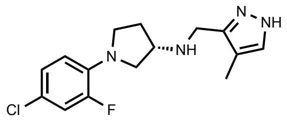 | 1.63 | 0.02 |
| ZINC000897222313 | 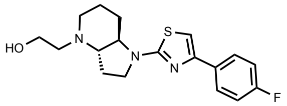 | 1.47 | 0.07 |
| ZINC000294308141 | 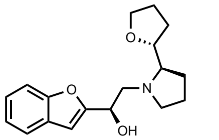 | 7.5 | 1.5 |
| ZINC000417864931 | 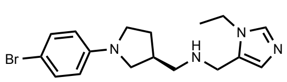 | 1.16 | 0.14 |
| ZINC000269761690 | 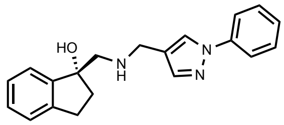 | 4.5 | 0.5 |
| ZINC000411862305 | 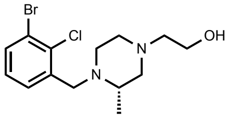 | 1.22 | 0.06 |
| ZINC000580533739 | 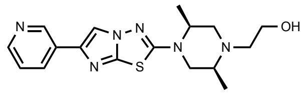 | 8 | 1 |
| ZINC000634713432 | 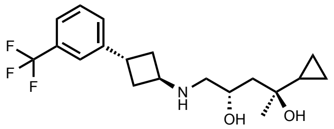 | 8.25 | 1.25 |
| ZINC000279319971 | 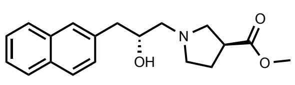 | 3 | 1 |
| ZINC000348114651 | 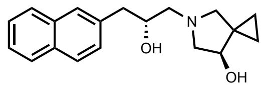 | 2.5 | 0.5 |

**Table S2: Active analogs of top five hits tested at SERT, related to Figure 1, S1 and STAR methods.**

| ZINC-ID/Supplier code | Comment | Chemical Structure | Ki, µM | SD |
|---|---|---|---|---|
| **ZINC000305339642** | **Initial hit** | 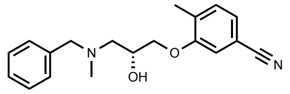 | **0.029** | **0.010** |
| ZINC000677296032 | analog of ZINC000305339642 | 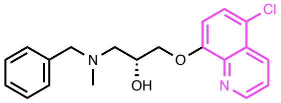 | 0.141 | 0.020 |
| ZINC000387330406 | analog of ZINC000305339642 | 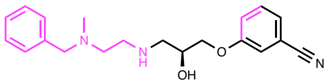 | 0.173 | 0.033 |
| Z4135312238 | analog of ZINC000305339642 | 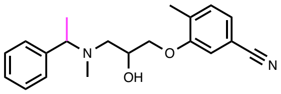 | 0.039 | 0.008 |
| Z4135312239 | analog of ZINC000305339642 | 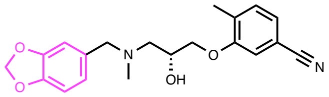 | 0.026 | 0.005 |
| **ZINC000623756919** | **Initial hit** | 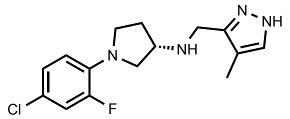 | **1.63** | **0.02** |
| ZINC000863582102 | analog of ZINC000623756919 | 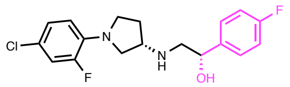 | 0.14 | 0.02 |
| ZINC000449730871 | analog of ZINC000623756919 | 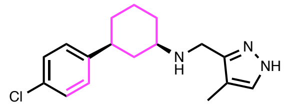 | 0.71 | 0.11 |
| Z4145515674 | analog of ZINC000623756919 | 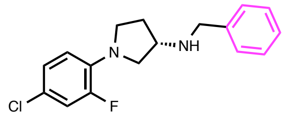 | 2.50 | 0.39 |
| Z4167665784 | analog of ZINC000623756919 | 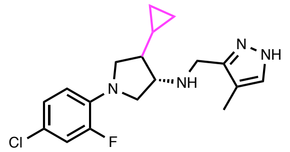 | 2.21 | 0.33 |
| **ZINC000897222313** | **Initial hit** | 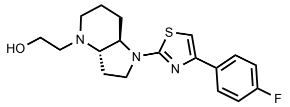 | **1.47** | **0.07** |
| ZINC000866205875 | analog of ZINC000897222313 | 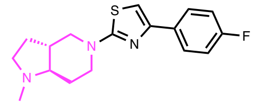 | 0.057 | 0.015 |
| ZINC000443438219 Racemic mix -8221 | analog of ZINC000897222313 | 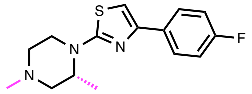 | 0.016 | 0.002 |
| ZINC00044343**8219** | R-isomer of ZINC000443438219 | 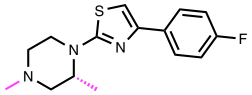 | 0.003 | 0.0004 |

| | | | | |
|---|---|---|---|---|
| ZINC000443438221 | S-isomer of ZINC000443438219 | 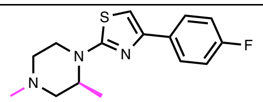 | 0.17 | 0.06 |
| ZINC000236204386 | analog of ZINC000443438219 | 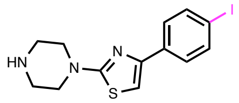 | 0.42 | 0.21 |
| ZINC000044494100 | analog of ZINC000443438219 | 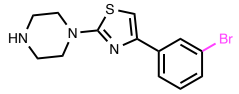 | 0.016 | 0.003 |
| ZINC000277296162 | analog of ZINC000897222313 | 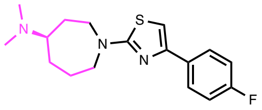 | 0.031 | 0.010 |
| ZINC000420267969 | analog of ZINC000897222313 | 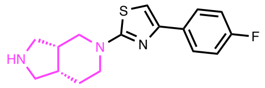 | 0.048 | 0.009 |
| ZINC00000665**8090** | analog of ZINC000897222313 | 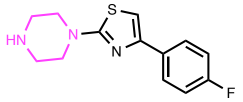 | 0.014 | 0.007 |
| ZINC000900380018 | analog of ZINC000897222313 | 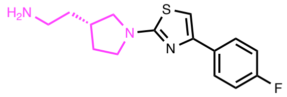 | 0.50 | 0.14 |
| **ZINC000417864931** | **Initial hit** | 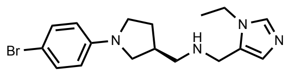 | **1.16** | **0.14** |
| ZINC000657329003 | analog of ZINC000417864931 | 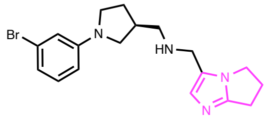 | 0.19 | 0.08 |
| ZINC000651685066 | analog of ZINC000417864931 | 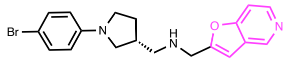 | 0.99 | 0.03 |
| ZINC000894225026 | analog of ZINC000417864931 | 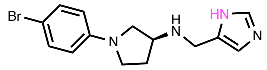 | 0.40 | 0.13 |
| **ZINC000411862305** | **Initial hit** | 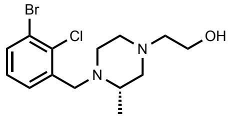 | **1.22** | **0.06** |
| ZINC001170690351 | analog of ZINC000411862305 | 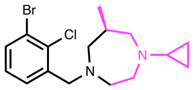 | 0.11 | 0.04 |
| ZINC001332938263 | analog of ZINC000411862305 | 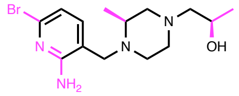 | 0.27 | 0.18 |
| Z4135312264 | analog of ZINC000411862305 | 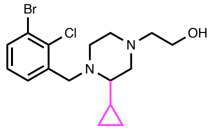 | 0.28 | 0.19 |

**Table S3**: **Cryo-EM data collection, refinement and validation statistics, related to Figure 5 and Figure S7.**

| | **SERT-Fab15B8 ‘8090-bound (EMDB: EMD-26160) (PDB: 7TXT)** |
|---|---|
| **Data collection** | |
| Microscope | Thermo Scientific Krios G3i |
| Detector | Gatan K3 with Gatan BioQuantum Energy filter |
| Voltage (kV) | 300 |
| Magnification | 105,000 |
| Defocus range (μm) | -1.0 to -2.1 |
| Pixel size (Å) | 0.81 (physical) |
| Total exposure ($e^-$/Å$^2$) | 50 |
| Frame exposure ($e^-$/Å$^2$/frame) | 0.833 |
| Images, number of | 5,010 |
| Frames/image, number of | 60 |
| Initial particles, number of | 3,313,742 |
| Final particles, number of | 187,696 |
| Symmetry imposed | C1 |
| Map sharpening *B* factor (Å$^2$) | -100 |
| Map resolution (Å) | 3.0 |
| FSC threshold | 0.143 |
| **Refinement and validation** | |
| Initial model used (PDB code) | 6VRH |
| Model resolution (Å) | 3.1 |
| FSC threshold | 0.5 |
| Model composition | |
| Chains | 4 |
| Non-hydrogen atoms | 6093 |
| Protein residues | 772 |
| Water | N/A |
| Ligands | 1 |
| *B* factors (Å$^2$) | |
| Protein | 43 |
| Ligand | 20 |
| R.m.s. deviations | |
| Bond length (Å) | 0.007 (1) |
| Bond angles (°) | 0.941 (0) |
| Validation | |

| | |
|---|---|
| MolProbity score | 1.56 |
| Clash score | 8.18 |
| EMRinger score | 4.50 |
| Rotamer outliers (%) | 0.00 |
| Ramachandran plot | |
| Favored (%) | 97.39 |
| Allowed (%) | 2.61 |
| Disallowed (%) | 0.00 |

**Table S4. Affinity differs between binding and transport assays, related to Figure 3 and STAR methods.**

| | '8219 | | '8090 | |
|---|---|---|---|---|
| | IC50 | SD | IC50 | SD |
| Binding | 0.0033 | 0.0008 | 0.0137 | 0.0032 |
| Transport, no preinc. | 0.113 | 0.023 | 0.280 | 0.053 |
| Transport, 30 m preinc. | 0.155 | 0.037 | 0.235 | 0.048 |

Binding measurements represent the IC50 values for displacement of [$^{125}$I]β-CIT from membranes prepared from SERT-expressing HeLa cells. Transport measurements represent IC50 values for inhibition of [$^{3}$H]5-HT transport into SERT-expressing HeLa cells with or without a 30 m preincubation with inhibitor.

**Table S5. Calculated binding and dissociation rates for '8219 and '8090, related to Figure 3 and STAR methods.**

| | **'8219** | **'8090** |
|---|---|---|
| $K_I$ – binding (M) | $5 \times 10^{-9}$ | $1.5 \times 10^{-8}$ |
| $k_{off}$ ($s^{-1}$) | $2.27 \times 10^{-3}$ | $1.82 \times 10^{-3}$ |
| $k_{on}$ ($M^{-1} s^{-1}$) | $4.54 \times 10^{5}$ | $1.21 \times 10^{5}$ |
| t½ at $K_I$, min | 5.09 | 6.35 |
| $K_I$ – transport (M) | $1.34 \times 10^{-7}$ | $2.58 \times 10^{-7}$ |
| $k_{on}$ ($M^{-1} s^{-1}$) | $1.69 \times 10^{4}$ | $7.07 \times 10^{3}$ |
| t½ at $K_I$, min | 0.19 | 0.37 |

For each compound, the rate of binding ($k_{on}$) was calculated from the measured dissociation rate ($k_{off}$) and the half-maximal inhibitory concentration ($K_I$) for transport and binding. The predicted half-time (t½) of the binding time course was then calculated at the $K_I$ concentration for each process. The half-times were much shorter than the 30 min. preincubation used in the experiment shown in Figure S4A, indicating that the difference in inhibitory potency in the two assays was not due to insufficient time for the compounds to equilibrate with SERT.

**Table S6. Dependence of affinity on $Na^+$ and $Cl^-$ ions, related to Figure 3 and STAR methods.**

| | **‘8219** | | **‘8090** | |
|---|---|---|---|---|
| | IC50 | SD | IC50 | SD |
| NaCl | 0.0040 | 0.0004 | 0.016 | 0.006 |
| Na-Isethionate | 0.037 | 0.011 | 0.023 | 0.003 |
| NMDG-Cl | 0.508 | 0.056 | 0.176 | 0.046 |

Binding affinity, as measured by the ability to displace [$^{125}$I]β-CIT from membranes prepared from SERT-expressing HeLa cells. Standard deviations were calculated from results of 3 separate experiments. For **‘8219**, the increased IC50 values in the absence of either $Cl^-$ or $Na^+$ was significant in a 2-sample t-test ($P<0.05$). For **‘8090**, the IC50 increase in the absence of $Cl^-$ was not significant ($P>0.05$) but the increase in the absence of $Na^+$ was significant ($P<0.05$).
